# Clinal genomic analysis reveals strong reproductive isolation across a steep habitat transition in stickleback fish

**DOI:** 10.1101/2020.08.28.269753

**Authors:** Quiterie Haenel, Krista B. Oke, Telma G. Laurentino, Andrew P. Hendry, Daniel Berner

## Abstract

How ecological divergence causes strong reproductive isolation between populations in close geographic contact remains poorly understood at the genomic level. We here study this question in a stickleback population pair adapted to contiguous, ecologically different lake and stream habitats. Dense clinal whole-genome sequence data reveal numerous regions fixed for alternative alleles over a distance of just a few hundred meters. This strong polygenic adaptive divergence must constitute a genome-wide barrier to gene flow because a steep cline in allele frequencies is observed across the entire genome, and because the cline center co-localizes with the habitat transition. Simulations confirm that such strong reproductive isolation can be maintained by polygenic selection despite high dispersal and small per-locus selection coefficients. Finally, comparing samples from the cline center before and after an unusual ecological perturbation demonstrates the fragility of the balance between gene flow and selection. Overall, our study highlights the efficacy of divergent selection in maintaining reproductive isolation without physical isolation, and the analytical power of studying speciation at a fine eco-geographic and genomic scale.

## Introduction

Deciphering the origin of species requires understanding the nature of reproductive isolation between diverging populations (Rice & Hostert 1993 ^1^; Coyne & Orr 2004 ^2^; Gavrilets 2004 ^3^; Sobel et al. 2010 ^4^). At the genomic level, reproductive isolation is typically studied through the marker-based comparison of population pairs that diverged relatively recently into ecologically different habitats (Lawniczak et al. 2010 ^5^; Ellegren et al. 2012 ^6^; Martin et al. 2013 ^7^; Toews et al. 2016 ^8^; Elgvin et al. 2017 ^9^). Genome regions showing exceptionally strong genetic population differentiation are inferred to harbor loci important to adaptive divergence that potentially also restrict the exchange of genetic material across larger chromosome segments, or the genome as a whole (Wu 2001 ^10^). This common analytical approach largely ignores the eco-geography of speciation. Our understanding of the genomics of reproductive isolation, however, can benefit greatly from investigating diverging populations across their contact zones at a fine geographic scale (Ryan et al. 2017 ^11^; Stankowski et al. 2017 ^12^; Pulido-Santacruz et al. 2018 ^13^; Rafati et al. 2018 ^14^; Westram et al. 2018 ^15^; Capblancq et al. 2020 ^16^). One reason is that such a clinal focus can reveal over what distance gene flow between populations occurs. Moreover, if marker resolution is sufficiently high – ideally whole-genome resolution, we can learn to what extent gene flow differs among genome regions. These details are crucial for evaluating the completeness and genetic architecture of reproductive isolation. Another benefit is that clinal analyses facilitate recognizing a possible link between reproductive isolation and ecological transitions, and hence the role of adaptation in initiating speciation (Schilthuizen 2000 ^17^; Sobel et al. 2010 ^4^).

Nevertheless, research combining a clinal perspective with the analytical power of whole-genome sequence data is largely lacking (Rafati et al. 2018 ^14^). Here, we present such an investigation in a threespine stickleback fish population pair residing in parapatriy (that is, contiguously) in Misty Lake and its inlet stream (Vancouver Island, British Columbia, Canada) (Lavin & McPhail 1993 ^18^; Hendry et al. 2002 ^19^) (Fig. 1a). This system is relatively young (postglacial, < 10,000 generations old) and the populations are fully genetically compatible when crossed (Lavin & McPhail 1993 ^18^; Hendry et al. 2002 ^19^; Raeymaekers et al. 2010 ^20^). Ecological differences between the lake and stream habitat, however, have driven dramatic genetically-based adaptive divergence in several traits including body shape, breeding coloration, trophic morphology and behavior (Lavin & McPhail 1993 ^18^; Hendry et al. 2002 ^19^; Sharpe et al. 2008 ^21^; Raeymaekers et al. 2009 ^22^; Berner et al. 2011 ^23^; Hanson et al. 2016 ^24^) (Fig. 1b, Fig. 2a, Supplementary Fig. 1). Low-resolution marker data further indicate that phenotypic divergence between the two populations is paralleled by substantial genetic differentiation (Moore et al. 2007 ^25^; Kaeuffer et al. 2012 ^26^; Stuart et al. 2017 ^27^). Given the absence of physical dispersal barriers separating Misty Lake from its inlet stream, and the potential of stickleback to disperse over hundreds of meters in a few days (even against water current; Bolnick et al. 2009 ^28^; Moore & Hendry 2009 ^29^), effective reproductive barriers between the populations must exist.

**Figure 1:**
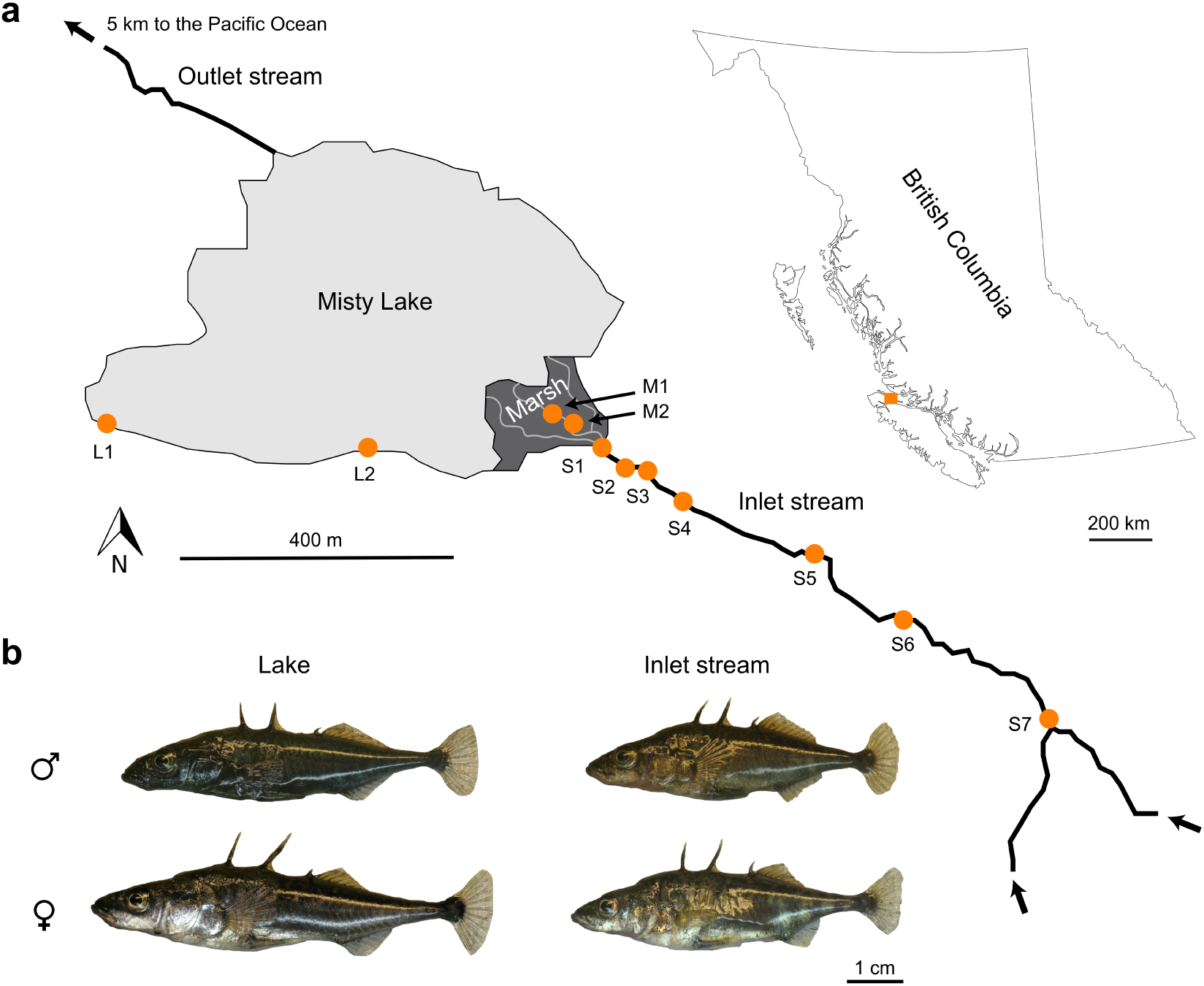
The Misty Lake and inlet stream stickleback system. (a) Geographic situation of Misty Lake, its inlet stream, and the marsh located between these habitats (map created by the authors based on data from Google Earth). Sample sites along the lake-stream transition are indicated by orange dots (GPS coordinates and sample sizes given in Supplementary Table 1). (b) Representative lake and stream stickleback individuals of both sexes (Photo credit: Katja Räsänen).

**Figure 2:**
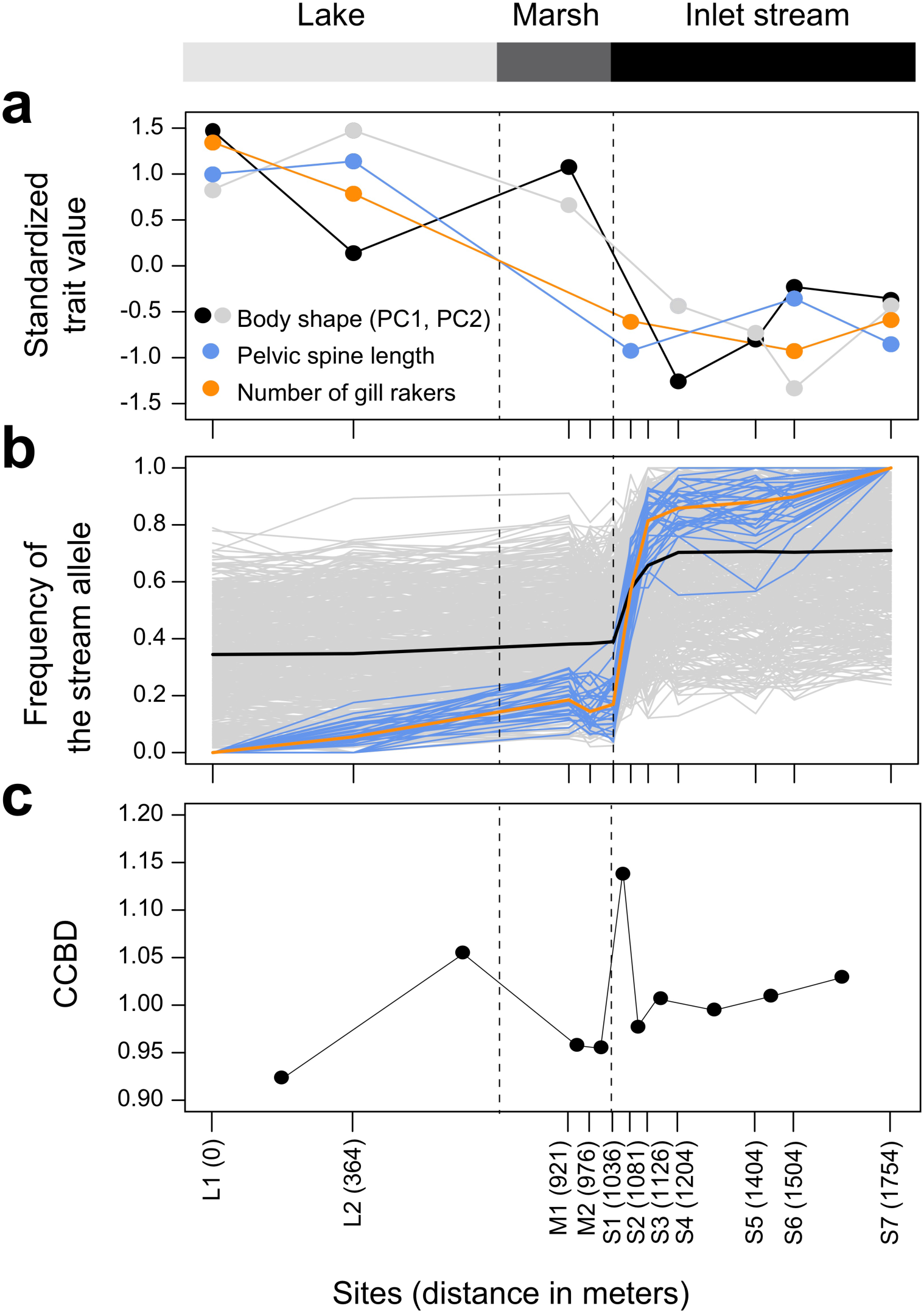
Phenotypic and genomic differentiation along the lake-stream transition. (a) Morphology, including geometric morphometric body shape (principal component scores, details in Supplementary Fig. 1), a predator defense trait (pelvic spine length), and a foraging trait (gill raker number). For ease of presentation, only site means are shown. Data were available for a subset of the study sites only. (b) Frequency of the stream allele at the 34 ‘fixed SNPs’ showing complete differentiation between the most distant sites L1 and S7 (blue lines; median frequency in orange), and at 500 random SNPs (gray lines; median frequency in black). (c) Chromosome center-biased differentiation (CCBD), calculated for all pairs of neighboring samples. As location along the gradient, the midpoint between the paired samples was used. In all panels, the vertical dotted lines indicate the lake-marsh and marsh-stream habitat boundaries. The distances on the x axis represent the approximate swimming distance between each site and the lake site L1.

To identify these reproductive barriers between Mistly lake and stream stickleback at the genomic level and to assess their impact on gene flow, we performed a clinal investigation based on pooled whole-genome sequencing at a fine geographic scale. Combined with individual-based simulations tailored to this system, our study offers a fine-grained illustration of how divergent natural selection can drive and maintain reproductive isolation between populations in direct contact.

## Results and Discussion

### Polygenic adaptive divergence between lake and stream stickleback

To initiate our investigation, we sampled approximately 56 stickleback individuals from each of 11 sites across the Misty Lake and inlet stream transition, with a distance of less than 2 km between the most distant sites (Fig. 1a). Each sample was subjected to pooled whole-genome sequencing to about 100x read depth, yielding approximately 1.9 million single-nucleotide polymorphisms (SNPs) as genetic markers.

We hypothesized that phenotypic and genetic divergence between the lake and stream fish maintained at a small geographic scale must be tightly linked to divergent selection between the habitats. Our first objective was therefore to identify genome regions likely targeted by selection by performing a standard pairwise population comparison. We here focused on our two most distant lake and stream sites (i.e., L1 versus S7, Fig. 1a), assuming that these represented our stickleback samples the least influenced by gene flow, and hence the best adapted to local lake or stream ecology. For this site pair, we quantified the magnitude of genetic differentiation, expressed by the absolute allele frequency difference AFD (Berner 2019 ^30^), across all genome-wide SNPs.

This genome scan revealed a median differentiation of 0.35 across all SNPs, thus confirming the strong overall genomic differentiation between the lake and stream population indicated previously by sparser marker data (Hendry et al. 2002 ^19^; Moore et al. 2007 ^25^; Kaeuffer et al. 2012 ^26^; Stuart et al. 2017 ^27^). However, numerous genomic regions exhibited much stronger differentiation. In particular, we identified 34 independent (i.e., separated by at least 100 kb) chromosome regions containing SNPs fixed for alternative alleles between the two most distant samples (differentiation along a representative chromosome is presented in Fig. 3, and along all chromosomes in Supplementary Fig. 2). For further analysis, one such SNP was chosen from each of these 34 regions, yielding our panel of ‘fixed SNPs’ representing putative genomic targets of strong divergent selection. Interrogating the gene annotation of the stickleback genome revealed that all 34 fixed SNPs lay outside coding gene sequences, consistent with primarily regulatory evolution during stickleback diversification (Jones et al. 2012 ^31^). Overall, our comparison of a single site pair makes clear that genomic divergence in Misty lake-stream stickleback is strong and highly polygenic; divergent selection likely targets hundreds of loci, as inferred in other lake-stream stickleback systems (Feulner et al. 2015 ^32^; Stuart et al. 2017 ^27^; Laurentino et al. 2020 ^33^).

**Figure 3:**
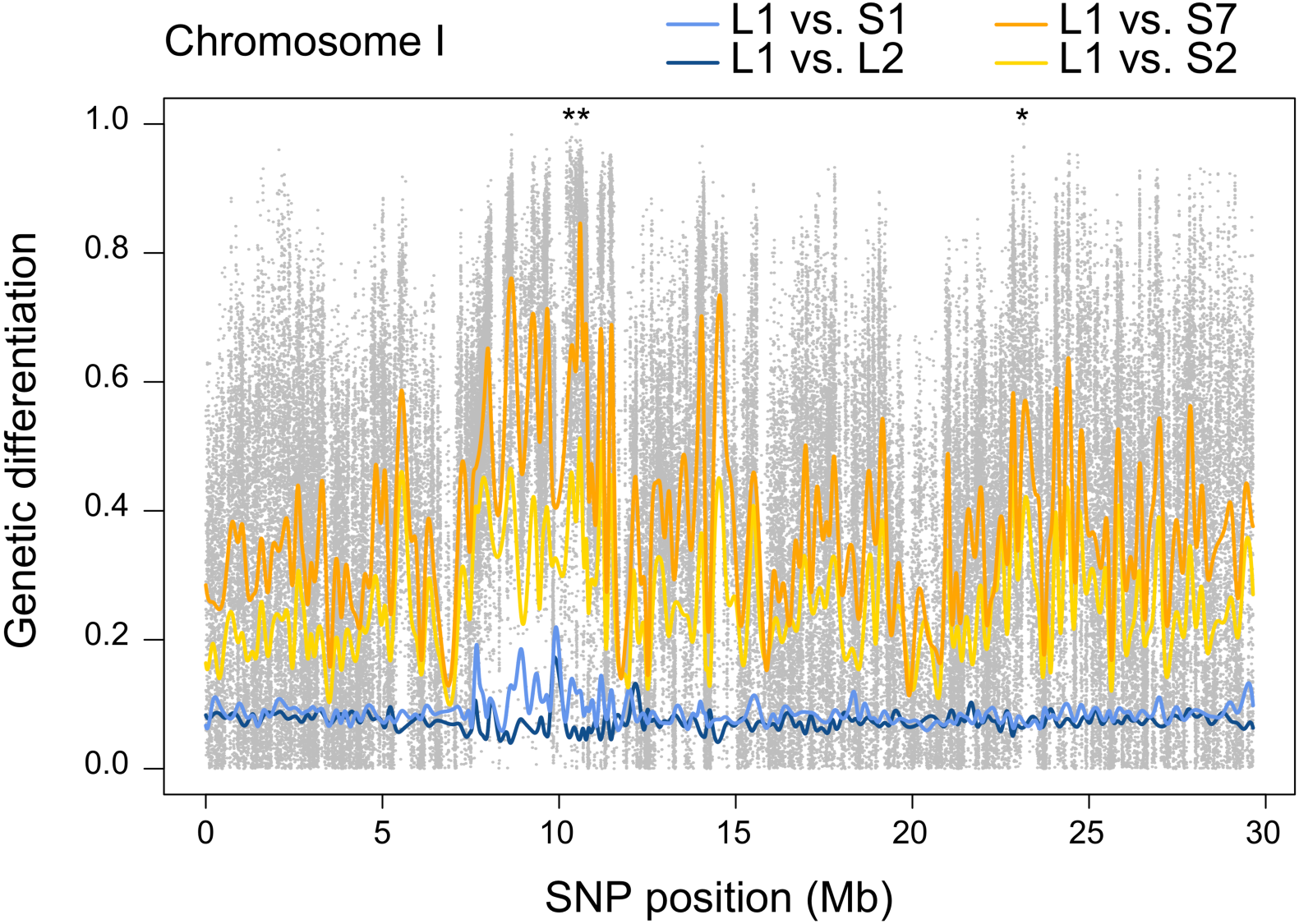
Pairwise genetic differentiation between clinal sites along a representative chromosome. Genetic differentiation is expressed as absolute allele frequency differentiation (AFD) for four site comparisons, each involving the lake site L1. The colored lines represent differentiation smoothed across SNPs by non-parametric regression. For the comparison between the endpoint samples (L1-S7), differentiation is additionally shown for all SNPs individually (gray dots), with fixed SNPs (AFD = 1) indicated by asterisks (*). Note that the differentiation profile from the L1-S1 comparison resembles the within-lake (L1-L2) comparison, except for a few genome regions particularly strongly differentiated in the L1-S1 comparison. The latter also prove exceptionally strongly differentiated between the lake and the more distant stream sites.

### Strong reproductive isolation at the habitat transition

Having obtained genomic evidence of the general presence of divergent selection, we took advantage of our full spatial set of samples to explore how tightly divergence was related to eco-geography. Graphing the frequency of the ‘stream allele’ (the allele fixed in the stream) at the fixed SNPs across all 11 clinal sites uncovered a dramatic genetic shift over a physical stream distance of merely 45 meters, its location coinciding with the transition of the marsh to the stream (blue and orange lines in Fig. 2b). This habitat transition also coincided with a major shift in ecologically important traits, as revealed by morphological data from this and previous studies for a subset of our sites (Fig. 2a). To expand our focus beyond adaptive genetic and phenotypic variation, we next inspected in all samples the frequency of the stream allele (here defined as the allele more frequent in sample S7 than L1) at 500 SNPs randomly chosen across the whole genome (the 34 fixed SNPs excluded). These ‘random’ SNPs also exhibited a steep cline coinciding with the marsh-stream transition (Fig. 2b, gray and black lines).

Together, these spatially fine-grained analyses reveal a remarkably tight association between ecology, phenotype, and genome-wide genetic variation. Thus, divergent selection not only maintains adaptive divergence, but simultaneously generates effective barriers to gene flow across the entire genome. What are these barriers? A major contribution to reproductive isolation must arise directly from local adaptation, in the form of reduced performance of migrants and hybrids between the habitats (Rice & Hostert 1993 ^1^; Coyne & Orr 2004 ^2^; Hendry 2004 ^34^; Nosil et al. 2005 ^35^; Sobel et al. 2010 ^4^). Field transplant experiments in a European lake-stream stickleback system indicate that these reproductive barriers alone can reduce lake-stream gene flow by around 80% (Moser et al. 2016 ^36^). Given that phenotypic differentiation in this European lake-stream system is relatively weak compared to other lake-stream systems (Berner et al. 2010 ^37^), selection against migrants and hybrids should cause an even stronger reproductive barrier in Misty stickleback. (In no parapatric lake-stream system assessed so far have crossing experiments under standardized laboratory conditions [Lavin & McPhail 1993 ^18^; Hendry et al. 2002 ^19^; Berner et al. 2011 ^23^; Eizaguirre et al. 2012 ^38^; Laurentino et al. 2020 ^33^] indicated hybrid unviability, thus ruling out intrinsic, ecologically-independent genomic incompatibility as an important reproductive barrier.) Additional reproductive barriers, like sexual isolation (Raeymaekers et al. 2010 ^20^; Berner et al. 2017 ^39^) or habitat preference (Bolnick et al. 2009 ^28^), likely contribute, but individually seem unlikely to cause the massive differentiation observed over small distances.

### The habitat transition is a narrow gene flow-selection tension zone

Having uncovered reproductive isolation across the habitat transition, we next asked how complete this isolation is. An informative pattern in our clinal genomic data was that the frequency of the stream allele at the random SNPs increased substantially upstream of the marsh-stream transition – but only up to sample site S4, that is, over c. 150 m (black curve in Fig. 2b). Beyond this location, allele frequencies proved stable over the entire remaining stream section investigated. The implication is that beyond a few hundred meters from the marsh-stream transition, reproductive isolation must already be nearly complete in the sense that any homogenizing effect of gene flow from the lake is no longer detectable. (This view does not conflict with the *fixed* SNPs showing some allele frequency change across the *entire* spatial gradient [orange curve in Fig. 2b]; the latter is expected from their particular ascertainment, i.e., complete differentiation between L1 and S7.)

Although stream allele frequencies at the random SNPs increased substantially over a few hundred meters upstream of the marsh-stream transition, these frequencies remained largely stable *downstream* of the transition, that is, all the way from site S1 across the marsh to the most distant lake site (black curve in Fig. 2b). This downstream stability indicates asymmetry in gene flow; genetic variation flows predominantly from the lake (and marsh; this habitat will be discussed in a later section) into the lower reach of the stream than in the opposite direction. While this asymmetry may be influenced by differences in dispersal behavior between lake and stream fish, we believe that a main reason is imbalance in relative population sizes: according to estimates from mark-recapture data, the lake population is substantially larger than the stream population (Oke et al. 2017 ^40^; Fisheries and Ocean Canada. 2018 ^41^). Moreover, contrary to the lake, the stream is a linear habitat; a relatively smaller fraction of the total stream population may have the opportunity to disperse into the lake than vice versa (see also Gavrilets 2004 ^3^).

### Gene flow is heterogeneous across the genome

The occurrence of gene flow from the lake to the lower reach of the stream allowed us to ask if some genome regions were more strongly reproductively isolated than others. A pattern relevant to this question emerged when quantifying genetic divergence by pairwise genomic comparisons of each of the sites L2 - S7 to the lake site L1, and graphing the resulting population differentiation along chromosomes. This complementary analysis revealed that the lowest stream site (S1; located right at the marsh-stream transition) was differentiated only trivially from the lake across most of the genome, hence producing chromosomal differentiation profiles closely resembling those from the within-lake (L1-L2) site comparison (compare the light-blue and dark-blue curves in Fig. 3 and Supplementary Fig. 2). Interestingly, however, a small number of genome regions exhibited substantially stronger differentiation in the L1-S1 comparison (e.g., around 10 Mb on chromosome I, Fig. 3). This pattern led us to hypothesize that our stream site closest to the lake (S1) was overwhelmed by gene flow from the lake, with appreciable genomic differentiation from the lake maintained only in the regions containing alleles most strongly selected in the stream. Under this hypothesis, we predicted that these regions should also exhibit exceptionally strong differentiation relative to randomly chosen genome regions when comparing the two samples from the endpoints of our geographic gradient (L1-S7). This prediction was confirmed by simulations estimating the magnitude of L1-S7 differentiation for the two classes of genome regions expected under drift alone (Fig. 4). Genome regions particularly highly differentiated between the lake and the closest stream site (S1) thus harbor loci under strong divergent selection that partly resist homogenization by gene flow.

**Figure 4:**
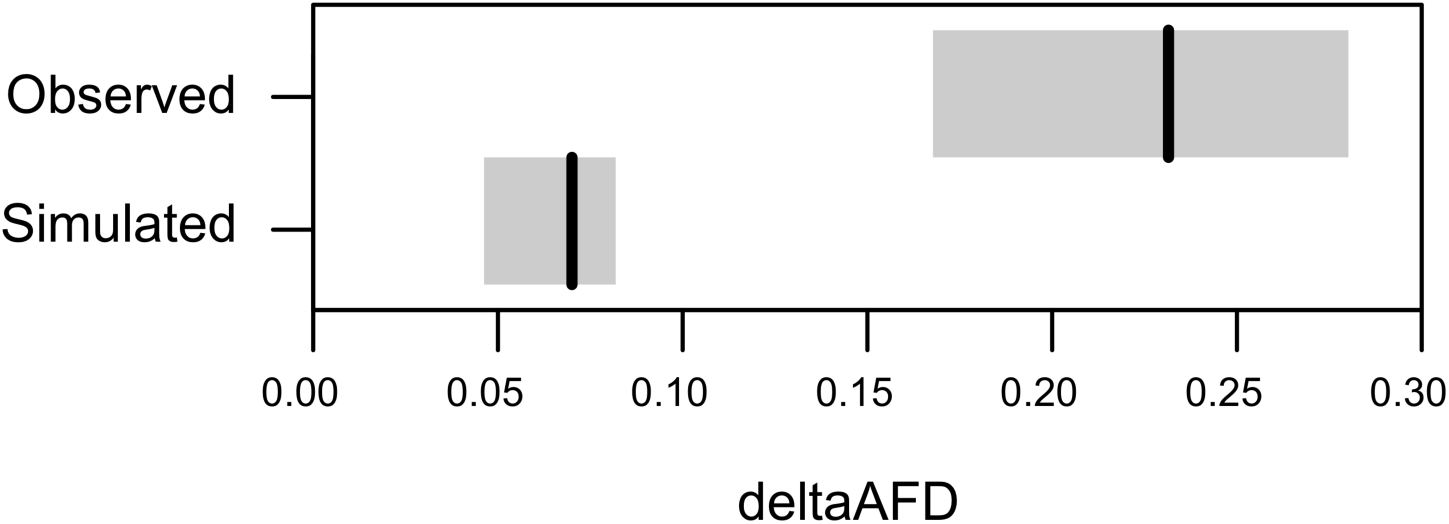
Divergent selection drives regions of high differentiation at the habitat transition. The upper row represents the empirically observed difference in AFD (deltaAFD) in the L1-S7 site comparison between high-differentiation regions and randomly chosen regions defined based on the L1-S1 comparison. The lower row represents deltaAFD calculated analogously for simulated high-differentiation and control loci evolving to similar baseline differentiation under drift alone. The gray boxes represent bootstrap 95% compatibility intervals. The deltaAFD observed empirically is much greater than expected under drift alone, indicating that the high-differentiation regions identified in the L1-S1 comparison must be under particularly strong divergent selection.

While this heterogeneity in gene flow between the populations concerned relatively small genome regions, we further obtained evidence of heterogeneous gene flow at the scale of whole chromosomes. Specifically, stickleback (as eukaryotes in general, Haenel et al. 2018 ^42^), exhibit substantially elevated crossover rates near the chromosome peripheries compared to the chromosome centers (Roesti et al. 2013 ^43^). Under this crossover distribution, theory predicts that polygenic divergent selection with gene flow causes relatively elevated population differentiation in chromosome centers (Roesti et al. 2012 ^44^; Berner & Roesti 2017 ^45^). We thus quantified the magnitude of chromosome center-bias in genetic differentiation (CCBD, Roesti et al 2012 ^44^, Berner & Roesti 2017 ^45^) for all pairwise combinations of neighboring samples, and found that this bias was greatest for the S1-S2 sample comparison (Fig. 2c). This supports the notion of an exceptionally strong antagonism between selection and gene flow in the lowest reach of the inlet stream.

Collectively, our analyses indicate that reproductive isolation between the Misty Lake and the inlet stream population *as a whole* is nearly complete, allowing the evolution of the two populations largely unconstrained by gene flow. Nevertheless, the lowest section of the stream represents a tension zone (Barton & Hewitt 1989 ^46^) in which selection is opposed by ongoing gene flow from the lake. This gene flow must involve hybridization between lake and stream fish, not just dispersal of lake fish into the lower stream section. The reason is that both heterogeneous genomic divergence in the L1-S1 sample comparison and the CCBD peak in the S1-S2 comparison require differential lake-stream gene flow among genomic regions, hence hybridization.

### Strong reproductive isolation in simulations of polygenic divergence

We have inferred from empirical patterns that reproductive isolation between parapatric stickleback populations reflects a by-product of adaptive divergence. To support the plausibility of our interpretation theoretically, we tailored individual-based simulations to the Misty stickleback system. We assumed nine demes in a linear array, with dispersal occurring in the beginning of every generation between contiguous demes only (stepping-stone model; Fig. 5a). Considering empirical population size estimates (Oke et al. 2017 ^40^; Fisheries and Ocean Canada. 2018 ^41^), the first (lake) deme was specified to be larger than all other (stream) demes together. The two habitats imposed polygenic divergent selection, local fitness being a function of genetic variation at 100 loci. All loci were polymorphic in the beginning of the simulations, thus mimicking standing genetic variation well known to underlie adaptive diversification in stickleback (Hohenlohe et al. 2010 ^47^; Deagle et al. 2013 ^48^; Roesti et al 2014 ^49^; Roesti et al. 2015 ^50^; Terekhanova et al. 2014 ^51^; Nelson & Cresko 2018 ^52^; Haenel et al. 2019 ^53^). After 1000 generations of evolution, we examined clinal patterns in the frequency of the stream allele (analogously to our empirical analyses of the fixed SNPs) resulting from different combinations of dispersal rates between the demes and strengths of divergent selection between the habitats.

**Figure 5:**
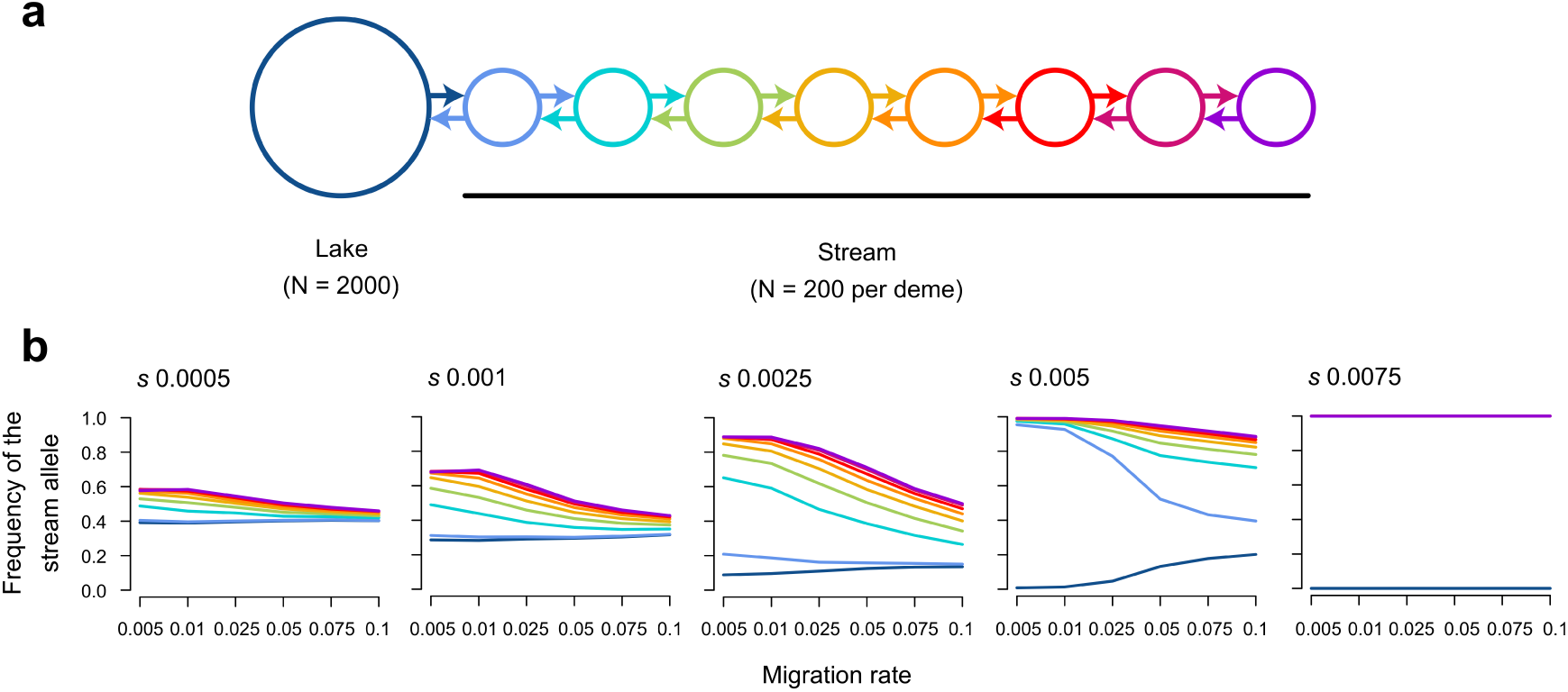
Simulated divergence with gene flow across a lake-stream habitat transition. (a) Schematic of the stepping stone model. Arrows indicate migration between neighboring demes. (b) Median frequency of the allele favored in the stream across 100 loci under selection after 1000 generations of evolution, for different combinations of migration rate and selection strength (selection coefficient *s* on top of each panel). The demes are color coded as in (a). The data are averages across 20 simulation replications. Note that the scaling of the x axis is not linear.

A first observation from these simulations was that strong adaptive divergence (and hence the associated reproductive barrier) established easily across the habitat transition, even for combinations of relatively small selection coefficients and high dispersal rates (Fig. 5b). Specifically, selection coefficients just below 0.01 already sufficed to allow complete differentiation between the lake and all stream sites across all dispersal rates considered (up to 0.1). Second, we found that whenever gene flow prevented complete lake-stream divergence, the stream site closest to the lake was particularly strongly constrained by gene flow, whereas more distant stream sites showed relatively similar allele frequencies (e.g., Fig. 5b, selection coefficients of 0.0025 or 0.005 combined with relatively high dispersal rates). This pattern closely resembled the shape of the cline in allele frequencies observed upstream of the marsh-stream transition at both our fixed and random SNPs (Fig. 2b). All these observations remained qualitatively consistent across our robustness checks (Supplementary Fig. 3).

Our simulations support our empirically-based conclusion that adaptive divergence from abundant standing genetic variation has produced strong reproductive isolation in the absence of physical barriers. Our study thus provides insights into the genomic architecture of adaptive divergence: previous theory has suggested that in the face of gene flow, adaptive divergence is promoted by the physical linkage of adaptive alleles, as produced by inversions (Rieseberg 2001 ^54^; Kirckpatrick and Barton 2006 ^55^; Yeaman 2013 ^56^). However, in Misty lake-stream stickleback, we found no indication of divergence in inversion polymorphisms. Importantly, the three large inversions often involved in adaptive divergence between pelagic and benthic stickleback ecotypes (Hohenlohe et al. 2010 ^47^; Jones et al. 2012 ^31^; Terekhanova et al. 2014 ^51^; Roesti et al. 2014 ^49^; Roesti et al. 2015 ^50^) proved monomorphic across our samples. Without denying the importance of chromosomal rearrangements in adaptive divergence in some study organisms, our whole-genome clinal investigation highlights that polygenic selection *per se*, without any particular physical arrangement of the targeted loci, can be sufficient for the emergence of strong divergence and reproductive isolation in the face of gene flow.

### Perturbation of gene flow-selection balance by an unusual ecological event

Our clinal phenotypic data and allele frequencies at the random SNPs revealed that the stickleback inhabiting the marsh (sites M1 and M2) are genetically very similar to the true lake samples. Nevertheless, in a previous study, microsatellite loci designed to discriminate Misty Lake and inlet stream fish indicated hybridization in the marsh (Hanson et al. 2016 ^24^). Similarly, the frequency of the stream allele at our fixed SNPs was slightly elevated in the marsh relative to the lake samples (although this may be attributable to the ascertainment of these SNPs, see above). This raised the possibility that the marsh might allow a modest degree of genetic mixing between the lake and the stream population despite being strongly dominated by lake fish. If true, we hypothesized that changes in the level of dispersal from the lake or the stream into the marsh, as mediated by a physical perturbation of the system, should drive a measurable shift in the genetic composition of the marsh fish.

Evaluating this hypothesis was made possible by exceptionally intense rainfall during our main sampling period, causing a rise in inlet stream discharge and lake water level that for a few days flooded the marsh normally above water level (Supplementary Fig. 4). To assess the genomic consequences of this event, we complemented our standard marsh sample (M1) taken before the flood by two additional samples from the same site, taken during the flood and one year later. The comparison of these temporal samples revealed a striking decline (often to zero) of the stream allele frequency at lake-stream population-distinctive SNPs during the flood, that is, within a few days (Fig. 6). Although our pooled sequence data precluded inspecting haplotype structure, the speed of this genetic change clearly indicates extensive dispersal of lake fish into the marsh, facilitated by easier access to the latter habitat. This conclusion is supported by simulations suggesting that at least 90% of the stickleback residing at the marsh site before the flood must have been replaced by lake immigrants during the flood (Supplementary Fig. 5). Interestingly, this perturbation in allele frequencies at the marsh site was partly offset one year later (Fig. 6), indicating selection on stream alleles and/or the new immigration of stream fish.

**Figure 6:**
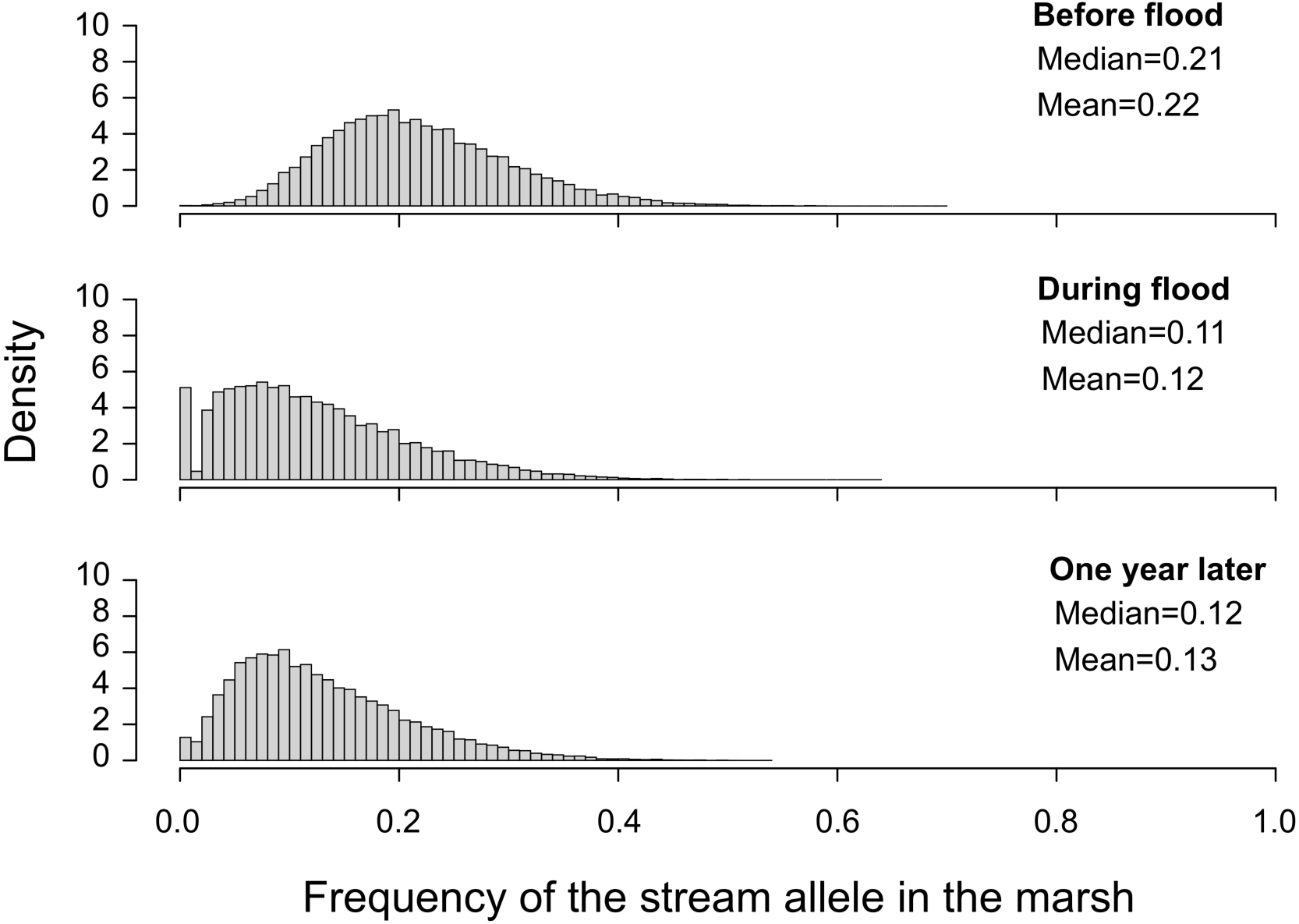
Impact of a flood on the genetic composition of stickleback in the marsh. Shown is the distribution of the frequency of the stream allele across 49,677 SNPs exhibiting strong lake-stream differentiation in stickleback sampled at the marsh site M1 at three different time points. The median and mean of the distributions are also given.

Our genomic analysis of temporally replicated samples from the marsh supports the idea that stickleback in this habitat represent a genomic mix between the lake and stream population caused by hybridization (Hanson et al. 2016 ^24^). Nevertheless, genetic material from the lake population vastly predominates, and we demonstrate that this imbalance can become nearly complete temporarily by an unusual short-term ecological modification of dispersal opportunities. Further disentangling the relative importance of selection and gene flow as determinants of allele frequencies within this eco-geographically intermediate habitat will benefit from direct information on the local selective conditions.

### Conclusions

Our investigation of parapatric stickleback has demonstrated the tight link between ecology, polygenic adaptive divergence, and whole-genome reproductive isolation in unprecedented detail, thereby illustrating how adaptation and speciation can be two sides of the same coin. Genetic exchange between the diverging populations, however, has not ceased completely but continues within a narrow tension zone. We show that the balance between homogenizing gene flow and divergent selection in this zone is fragile and can shift quickly when habitats are perturbed. Our work highlights the power of whole-genome sequencing at a fine spatial scale and across multiple time points to inform the eco-geography and genetic architecture of speciation.

## Methods

### Stickleback sampling and phenotypic analysis

Stickleback were captured with unbaited minnow traps at 11 sites in Misty Lake and its inlet stream between May and July 2016 (the marsh site M1 was additionally sampled in August 2017). Sample sizes ranged from 40-62 individuals per site (details on the locations and samples given in Supplementary Table 1). From each individual, a dorsal spine was clipped and stored in 95% ethanol, for DNA extraction. All individuals were immediately released.

To explore differentiation in geometric morphometric body shape along the geographic gradient, 40 individuals from a subset of our sites (L1, L2, M1, S4, S5, S6 and S7) were photographed on their left side with a standard scale using a digital camera (Canon PowerShot G11, Canon, Tokyo, Japan). All photographs were digitized with tpsDIG2 (life.bio.sunysb.edu/morph/) by the same investigator (KBO) in haphazard order by placing 14 landmarks used in previous studies in the same system (Kaeuffer et al. 2012 ^26^; Oke et al 2016 ^57^). Using the *geomorph* R package (Adams & Otarola-Castillo 2013 ^58^, R Core Team 2019 ^59^), the resulting coordinates were aligned and generalized Procrustes analysis performed, yielding principal components of body shape variation among individuals (Rohlf & Slice 1990 ^60^; Adams & Otarola-Castillo 2013 ^58^). In addition, we retrieved data for two ecologically important traits related to predator defense (pelvic spine length) and foraging (number of gill rakers) from another subset of our sites (L1, L2, S2, S6 and S7) studied in a previous phenotypic study of Misty Lake and inlet stickleback (Moore & Hendry 2005 ^61^). All these phenotypic data were mean-centered and standardized to allow visualization on the same scale.

### DNA library preparation and sequencing

DNA was extracted individually from the dorsal spine clip of each of the 701 total stickleback using the Quick-DNA TM Miniprep Plus Kit (Zymo Research, Irvine, CA, USA), by following the manufacturer protocol. For enzymatic tissue digestion, the spines were minced with micro spring scissors to maximize DNA yield. Following DNA quantification using a Qubit fluorometer (Invitrogen, Thermo Fisher Scientific, Wilmington, DE, USA), individuals were pooled to equal molarity without PCR-enrichment to obtain a single DNA library per sampling site (and per time point in the case of the marsh site M1). The 13 total libraries were paired-end sequenced to 151 base pairs on ten total lanes of an Illumina HighSeq2500 instrument, producing a median read depth per base pair of 103x across the samples (min=51, max=133; Supplementary Table 1). Combined with the relatively large number of individuals per site, this high read depth was expected to allow estimating allele frequencies with high precision (Ferretti et al. 2013 ^62^; Gautier et al. 2013 ^63^).

### SNP discovery

Raw sequences reads were parsed by sampling site and aligned to the third-generation assembly (Glazer et al. 2015 ^64^) of the stickleback reference genome (Jones et al. 2012 ^31^) by using Novoalign (Version 3.03.00, http://www.novocraft.com/products/novoalign/). The Rsamtools R package (Morgan et al. 2017 ^65^, R Core Team 2019 ^59^) was then used to convert the alignments to BAM format, and to perform nucleotide counts for all genome-wide positions using the *pileup* R function (Morgan et al. 2017 ^65^; R Core Team 2019 ^59^). To identify informative single-nucleotide polymorphisms (SNPs), we first combined nucleotide counts from the two lake samples (L1, L2) and from the two most upstream inlet samples (S6, S7) into lake and stream pools. Genomic positions then qualified as SNPs for further analysis if they exhibited a read depth between 50x and 400x within each pool, and a minor allele frequency (MAF) of at least to 0.25 across the two pools combined. The 1,920,596 SNPs passing these filters were genotyped in all 13 samples separately.

### Quantifying clinal genomic differentiation

Genetic differentiation across the lake-stream transition was quantified by two approaches. The first relied on the frequency of the stream allele at ‘fixed’ and ‘random SNPs’. Specifically, we screened our SNP panel for markers showing complete genetic differentiation, as quantified by the absolute allele frequency difference AFD (Berner 2019 ^30^; the differentiation metric used throughout our study), between the most distant samples (L1, S7). If multiple such SNPs were less than 100 kb apart, they were considered a cluster from which only one SNP was chosen at random. This produced a panel of 34 independent fixed SNPs. As a resource for future investigations, we retrieved from the reference genome annotation all genes located within a 100 kb window centered at each of the fixed SNPs, producing a gene compilation (provided as Supplementary Table 2) likely containing numerous genes under divergent lake-stream selection in the Misty system. The annotation was also used to assess if the fixed SNPs were located within or outside coding gene sequences. The random SNPs, in turn, represented 500 markers chosen at random across the genome, again applying a 100 kb spacing threshold. For each class of SNPs (fixed, random), we then identified the ‘stream allele’ as the nucleotide relatively more frequent (or fixed) in the S7 than the L1 sample. Finally, the frequency of the stream allele was calculated for each sample and visualized along the geographic gradient. In the second approach, we calculated genetic differentiation at all genome-wide SNPs for each pairwise combination of the first lake sample (L1) and all other samples. The values obtained were visualized along chromosomes, raw and/or smoothed by non-parametric regression using the *smooth*.*spline* R function (band width 0.1).

### Inferring selection-gene flow antagonism from high-differentiation regions

While inspecting the comparison L1-S1, we observed that some genome regions showed remarkably strong genetic differentiation compared to the overall undifferentiated genome-wide background. This led us to speculate that in these regions, genetic variants particularly strongly selected in the stream were maintained at elevated frequency at the S1 site, while the remainder of the genome was overwhelmed by gene flow from the lake. We thus predicted that these specific genome regions should also exhibit exceptionally strong differentiation in the comparison L1-S7. To evaluate this idea, we first subtracted the mean AFD value in the L1-L2 comparison (considered the differentiation baseline within the lake habitat) from the corresponding value in the L1-S1 comparison for each genome-wide 10 kb sliding window (5 kb overlap) containing at least 5 SNPs. For the most highly differentiated 0.5% of these windows (HDW; N = 178), considered regions under strong lake-stream selection, and for the same number of windows chosen at random as control (non-HDW), we then calculated mean AFD in the L1-S7 comparison.

Because the HDW by definition exhibited elevated differentiation in the L1-S1 comparison, somewhat stronger L1-S7 differentiation in these windows relative to random windows was expected even if the HDW were strongly differentiated in the former comparison just by chance. Comparing the two classes of windows thus required a benchmark, which was obtained by individual-based simulation. We here constructed *n* haploid individuals by first generating 178 biallelic (1, 0) loci (non-HDL) at which the stream allele (1) occurred at a frequency specified by a random draw from the uniform distribution bounded between 0.05 and 0.5 (i.e., the stream allele was always the minor allele, as observed empirically at the site S1; Fig. 2b). Another set of loci (HDL) was generated analogously, except that the frequency of the stream allele was increased by 0.1, corresponding to the observed difference in median L1-S1 AFD between the HDW and non-HDW. We then allowed this population to evolve neutrally (i.e., to drift) by drawing offspring for each new generation at random with replacement. All loci were unlinked, and random assortment of alleles was achieved by swapping alleles between the haplotypes within pairs of offspring. After *g* generations, we determined median AFD before vs. after evolution for the HDL and non-HDL. The combination of *n* and *g* was chosen to produce drift at the non-HDL approximating the AFD observed at the non-HDW in the L1-S7 comparison (*n* =200, *g* = 1000; higher values for both variables produced similar results but required longer simulation). This simulation was replicated 25 times.

Finally, we calculated the difference in median AFD in the L1-S7 comparison between the empirical HDW and non-HDW, and the median difference in AFD achieved during simulated evolution at the HDL and non-HDL across the 25 replicates, both referred to as deltaAFD. Uncertainty around these point estimates was obtained by bootstrapping windows (empirical data) and replicates (simulated data) 10,000 times. This allowed evaluating if differentiation at the HDW was consistent with drift alone, or indicated deterministic selection.

### Inferring selection-gene flow antagonism from CCBD

The combination of polygenic selection, gene flow, and a reduced crossover rate in chromosome centers compared to chromosome peripheries causes relatively elevated population differentiation in chromosome centers (chromosome center-biased differentiation, CCBD; Roesti et al. 2012 ^44^; Roesti et al. 2013 ^43^; Berner & Roesti 2017 ^45^; Haenel et al. 2018 ^42^). To explore the strength of selection-gene flow antagonism along the geographic gradient, we thus quantified to what extent genetic differentiation was biased toward chromosome centers for all ten pairwise comparisons of neighboring samples. For this, we defined the outer 5 Mb on either side of a chromosome as high-crossover rate periphery and the remainder of the chromosome as low-crossover rate center (Roesti et al. 2013 ^43^). Next, we divided median AFD across all central SNPs by median AFD across the peripheral SNPs for each chromosome within each site pair. For each pair, we then treated the median across these ratios as CCBD, and graphed this metric along the lake-stream gradient by using the midpoint between the neighboring samples as geographic location. As a robustness check, this analysis was repeated by using as site pairs each of the samples L2-S7 combined with L1, which produced very similar results supporting the same conclusion (Supplementary Fig. 6).

### Individual-based simulations of divergence with gene flow

To explore theoretically how selection can drive and maintain reproductive isolation in the presence of gene flow, we conducted individual-based forward simulations using a diploid stepping stone expansion of the model in Berner & Roesti 2017 ^45^. Our standard model involved nine total demes arranged in a one-dimensional array, with adjacent demes connected by migration (Fig. 5a). The first deme (N=2000) represented the (larger) lake population while all other demes (each N=200) represented stream sites. In the beginning of each generation, a fraction of *m*/2 individuals was chosen at random from each deme to migrate into the neighboring deme on either side (juvenile migration). Each deme thus had a total fraction of *m* individuals replaced by immigrants, except for the demes located at the endpoints of the array, for which this fraction was *m*/2. The migration phase was followed by selection and reproduction. We modeled polygenic divergent selection by assuming a total of 100 biallelic unlinked loci under divergent selection, with one allele favored in the first deme and the other allele favored in all other demes. The simulations started with the frequency of both alleles set to 0.5 in all demes, to minimize the probability of the stochastic loss of adaptive variation (Berner & Roesti 2017 ^45^). Loci contributed additively to fitness; each maladaptive allele reduced an individual’s fitness from the local fitness optimum of one by *s*, the selection coefficient. Individual fitness was then scaled by the mean population fitness and determined an individual’s probability to be drawn for reproduction. Individuals reproduced as hermaphrodites and were allowed to mate more than once, each mating producing one offspring. Mating was repeated until the number of offspring re-established local deme size. The offspring cohort then replaced the parental deme (discrete generations), and entered the migration phase.

We explored a range of combinations of migration rates (0.005 – 0.1) and selection coefficients (0.0005 - 0.0075). All simulations were run for 1000 generations, a time span shown by preliminary runs over up to 7000 generations to allow approaching migration-selection balance (Supplementary Fig. 7 a and b). All parameter combinations were replicated 20 times. Results were visualized by plotting for each deme at generation 1000 the mean across simulation replications of the median frequency of the allele favored in the stream across all 100 loci for all combinations of migration rates and selection coefficients.

The robustness of the standard model described above was assessed by a number of additional simulations. We here considered models with a multiplicative (as opposed to additive) contribution of each locus to overall fitness, a lower number of selected loci (10), physical linkage among all loci along a single chromosome undergoing crossover during mating (as opposed to independent segregation), and finally locus-specific selection coefficients drawn at random from the exponential distribution with a rate equal to 1/*s* (Supplementary Fig. 3).

### Quantifying the impact of a flood on stickleback in the marsh

For the marsh site (M1), three temporally replicated samples were available: before, during, and one year after a flood. The former represents the sample also used in all previous analyses; the latter two samples were processed in exactly the same way. To maximize the sensitivity for detecting gene flow, we here considered only SNPs highly differentiated (AFD >= 0.75) between the lake pool (sites L1 and L2 combined) and the stream pool (sites S6 and S7), and sequenced to a minimum read depth of 50x within each temporal sample. For the 49,677 SNPs thus obtained, we visually compared the frequency of the stream allele among the samples.

Because this analysis indicated massive dispersal of lake stickleback into the marsh during the flood, we explored by simulation how much of such dispersal was needed to drive the observed change in allele frequencies. For the SNPs above, we here combined allele frequency data from the marsh before the flood with those from the nearest lake site (L2), considering a wide range of different relative proportions of the latter (10% - 100%). Comparing the resulting (mixed) allele frequency distributions to the one observed during the flood allowed a qualitative assessment of the proportion of lake dispersers into the marsh during the flood.

Unless stated otherwise, all our analyses and simulations were implemented in the R language (R Core Team 2019 ^59^).

## Author contributions

D.B. initiated and supervised the study; D.B. and Q.H. designed the experiment; D.B., Q.H and A.P.H acquired funding; K.B.O. and A.P.H. performed field sampling and measurements; K.B.O. generated and analyzed phenotypic data; Q.H. and T.G.L. performed wet lab work; D.B. and Q.H. implemented analytical tools; Q.H. and D.B. analyzed genomic data and interpreted results; Q.H. and D.B. wrote the manuscript, with input from all co-authors.

## Acknowledgements

This project was supported financially by the Swiss National Science Foundation (grant 31003A_165826 to DB) and the Freiwillige Akademische Gesellschaft Basel (QH), and field sampling additionally by Fisheries and Oceans Canada, the British Columbia Ministry of the Environment, and the McGill University Biology Department. We thank Fiona Beaty, Brody Forst, Bailey Feddersen, Tristan Kosciuch, Minxin Lu, Erica MacClaren, Emily McIntosh, Alexanne Oke, Sarah Sanderson, and Mac Willing for aiding field sampling; Western Forest Products for providing logistical and safety support and access to field sites; Walter Salzburger for sharing web lab infrastructure; Brigitte Aeschbach and Nicolas Boileau for facilitating lab work; Christian Beisel, Ina Nissen and Elodie Burcklen for Illumina sequencing at the Quantitative Genomics Facility, D-BSSE, ETH Zürich; the developers of Novocraft for sharing their sequence aligner; Katja Räsänen for providing pictures of Misty stickleback; Laurent Guerard and Nicolás Lichilín Ortiz for help with scripting. Computation was performed at the sciCORE (https://scicore.unibas.ch) scientific computing center of the University of Basel.

## Data archiving

All raw Illumina sequences, demultiplexed by site (and sampling period for the site M1) are available from the European Nucleotide Archive, project PRJEB37366.

**Supplementary Information and Supplementary Codes are available for this paper**

