## Supplementary Information for "Clinal genomic analysis reveals strong reproductive isolation across a steep habitat transition in stickleback fish"

### Contents

#### Supplementary Figures

Supplementary Figure 1: Geometric morphometric analysis of lake, marsh and stream stickleback. (Page 3-4)
Supplementary Figure 2: Genetic differentiation between Misty Lake and inlet stream stickleback across all chromosomes. (Pages 5-7)
Supplementary Figure 3: Robustness checks of the simulations of divergence with gene flow across a habitat transition. (Page 8)
Supplementary Figure 4: Characterization of the marsh habitat in the Misty system. (Page 9)
Supplementary Figure 5: Exploring the approximate proportion of migrants from the lake into the marsh during the flood. (Page 10)
Supplementary Figure 6: Alternative analysis of chromosome center-biased differentiation (CCBD). (Page 11)
Supplementary Figure 7: Determining an appropriate number of generations for the simulations. (Page 12)

#### Supplementary Tables

Supplementary Table 1: Characterization of the study sites in the Misty Lake watershed. (Page 13)
Supplementary Table 2: List of all genes present within a 100 kb window around each of the 34 SNPs showing complete differentiation between the most distant lake and stream sites (L1, S7). (Pages 14-21)

#### References

#### Supplementary Figures

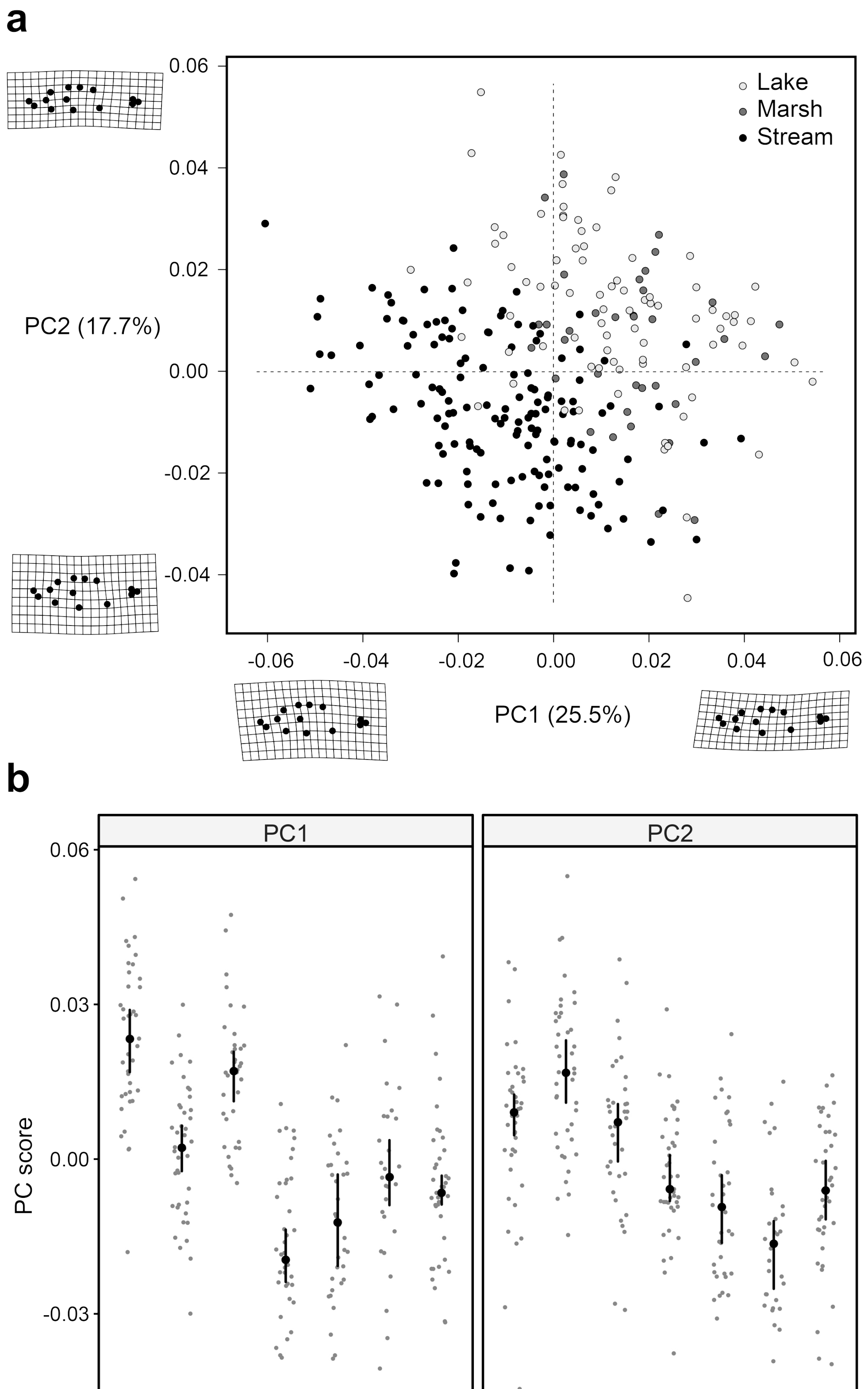

**Supplementary Figure 1**: **Geometric morphometric analysis of lake, marsh and stream stickleback.** (a) Each point represents an individual fish along the first two principal components (PC1, PC2) obtained by landmark-based shape analysis (methodological details given in Kaeuffer et al. 2012 and Oke et al. 2016). Individuals are color coded according to their habitat (lake, marsh, inlet stream). The deformation grids visualize the body shape associated with the lowest and highest observed score along each PC. (b) Individual PC scores shown separately for each study site. Black dots and vertical lines represent site medians with their bootstrap 95% compatibility interval. Note that both PCs capture variation in body depth, that stream fish tend to exhibit deeper bodies than lake fish, and that marsh fish resemble lake fish.

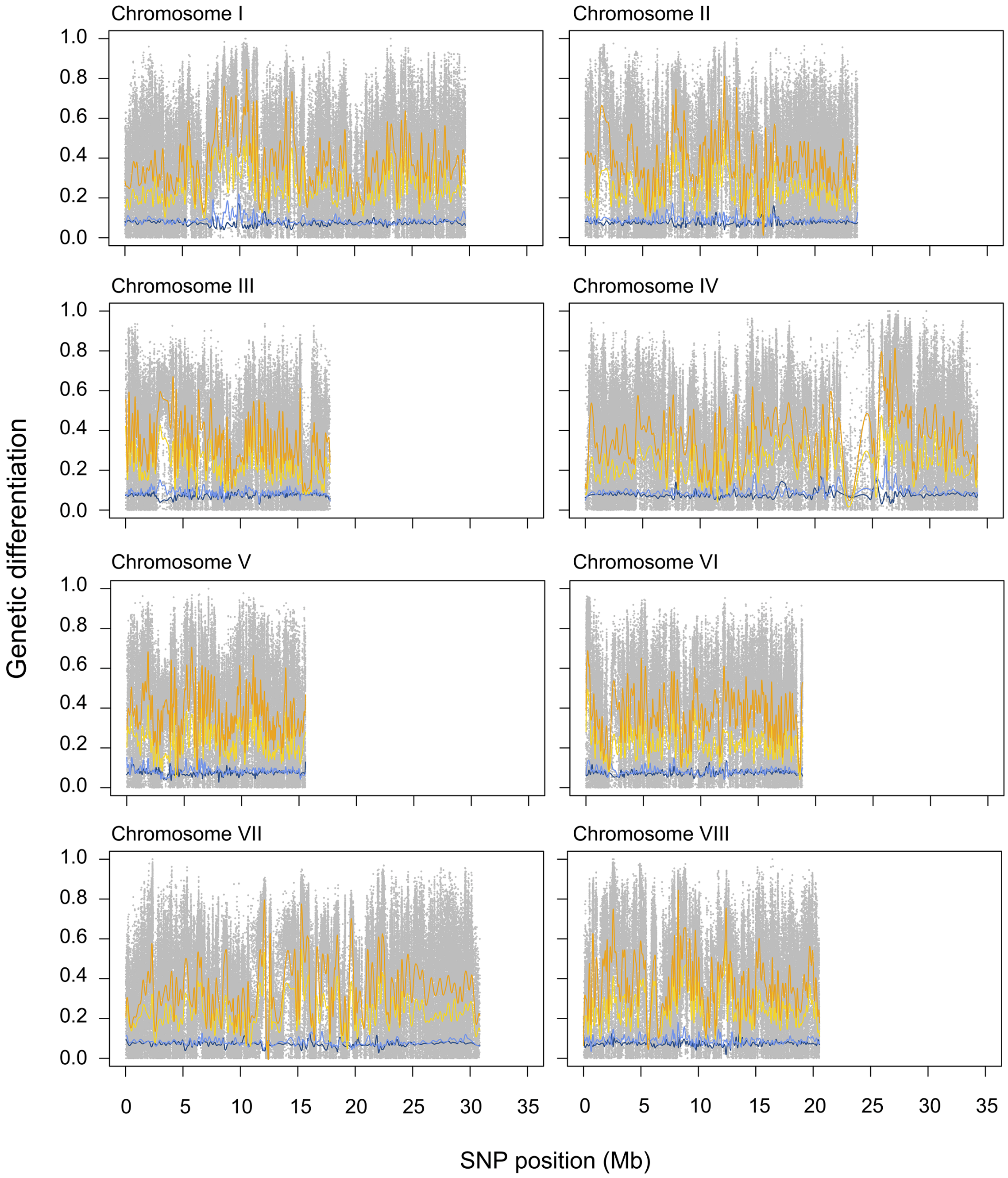

Chromosome I
Chromosome II
Chromosome III
Chromosome IV
Chromosome V
Chromosome VI
Chromosome VII
Chromosome VIII
1.0
0.8
0.6
0.4
0.2
0.0
0
5
10
15
20
25
30
35
Genetic differentiation
SNP position (Mb)

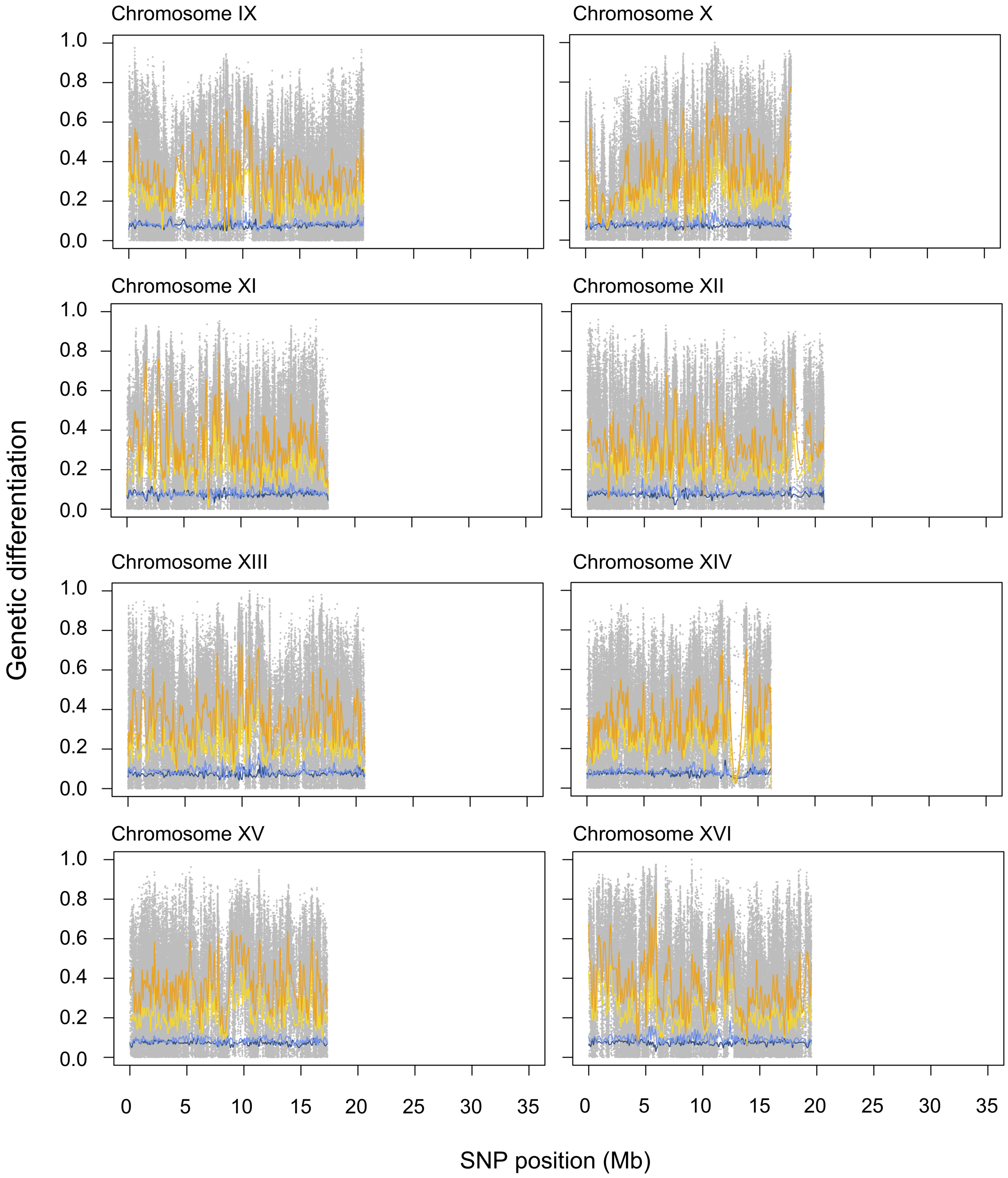
Chromosome IX
Chromosome X
Chromosome XI
Chromosome XII
Chromosome XIII
Chromosome XIV
Chromosome XV
Chromosome XVI
Genetic differentiation
1.0
0.8
0.6
0.4
0.2
0.0
0
5
10
15
20
25
30
35
SNP position (Mb)

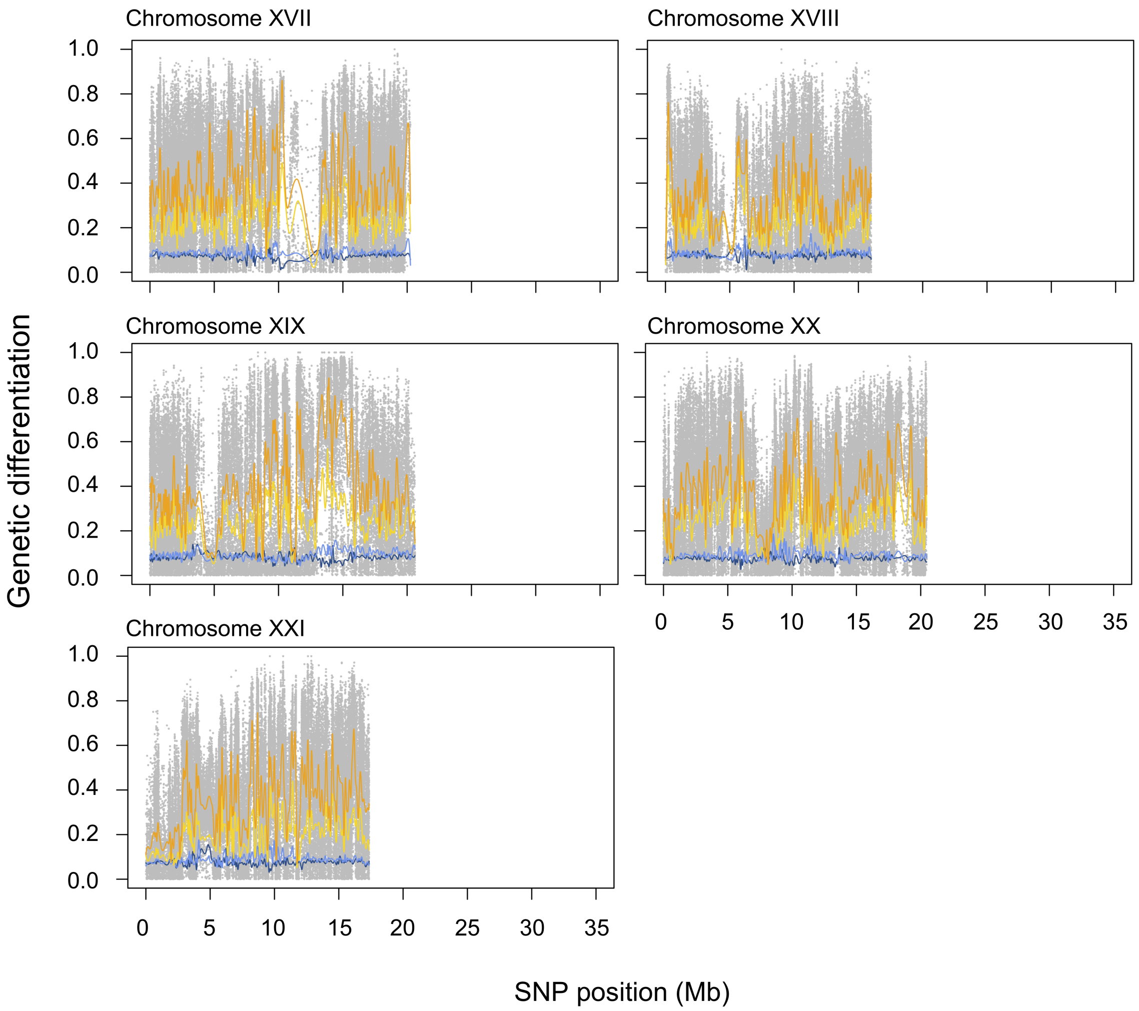

**Supplementary Figure 2: Genetic differentiation between Misty Lake and inlet stream stickleback across all chromosomes.** Differentiation is expressed by the absolute allele frequency difference AFD. The presentation format follows that of Fig. 3.

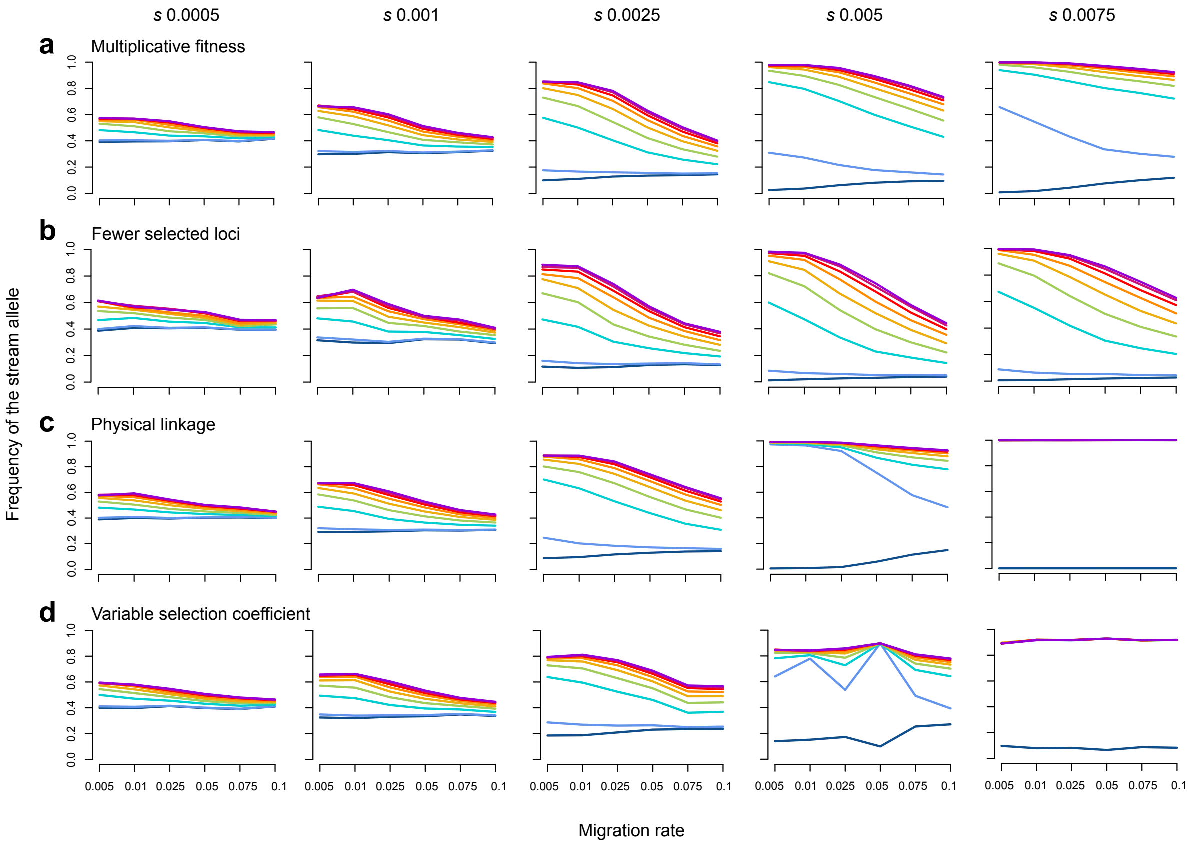

**Supplementary Figure 3: Robustness checks of the simulations of divergence with gene flow across a habitat transition.** The presentation format follows Fig. 5, but the simulations were performed with the standard model modified in the following ways: (a) The loci contribute to fitness multiplicatively, as opposed to additively. (b) Only ten loci are under selection, as opposed to 100. (c) All loci are physically linked on a single chromosome exhibiting crossover, as opposed to free segregation. (d) The selection coefficients are not identical among loci, but are drawn from an exponential distribution (rate = 1/*s*). Note that the latter causes greater stochasticity in the simulation outcome, as particularly evident with s=0.005.

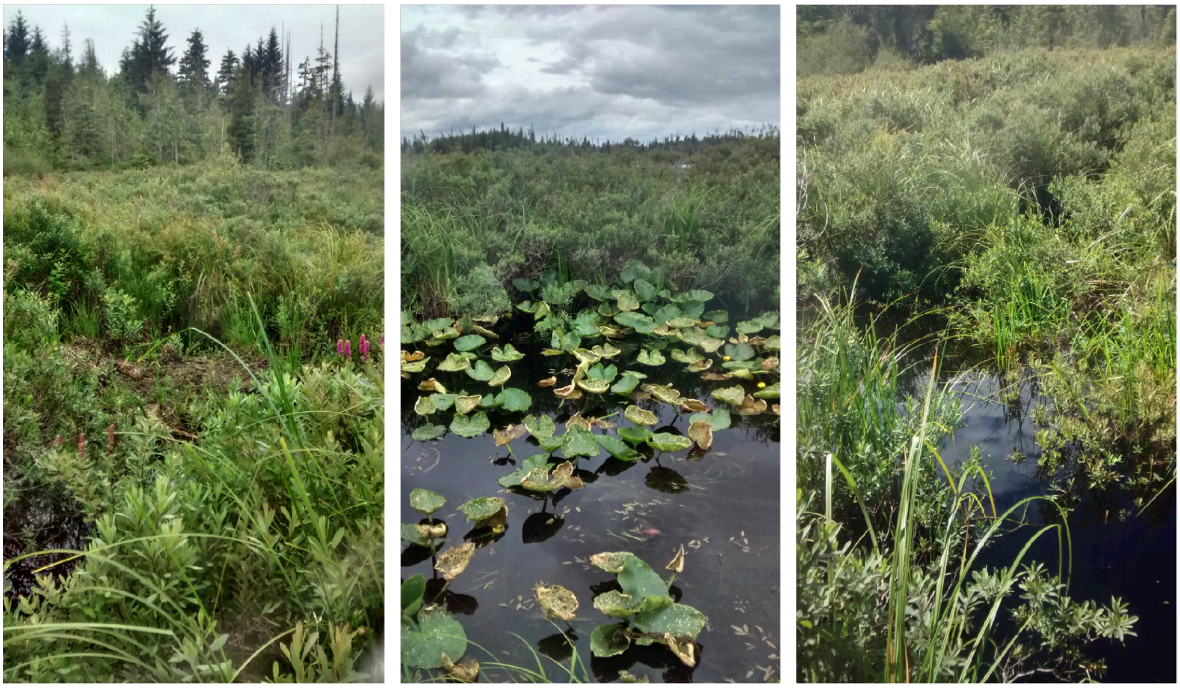

**Supplementary Figure 4: Characterization of the marsh habitat in the Misty system.** The marsh represents the transition from the inlet stream (flowing through woodland) to the lake. This habitat is dominated by short (<1.5 m), dense vegetation intersected by relatively narrow, deep channels. During most of the summer, one can walk on mats of this vegetation, the roots of which reach approximately 30 cm below the water surface. During the flood, these mats became entirely submerged, likely allowing fish to swim through vegetation that days before would have been above water. Supporting this view, stickleback catch rates at the marsh site increased dramatically during the flood (Krista B. Oke, unpublished data) (Photo credits: Krista B. Oke).

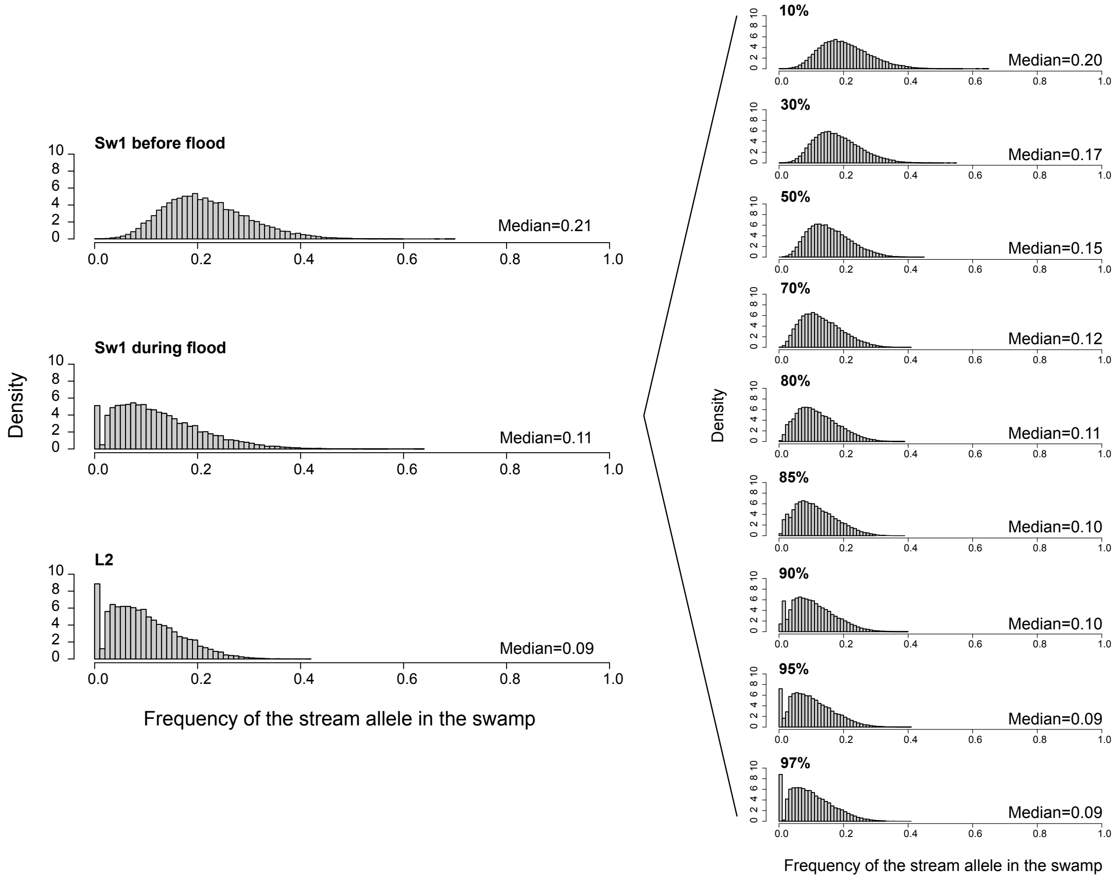

**Supplementary Figure 5: Exploring the approximate proportion of migrants from the lake into the marsh during the flood.** This analysis focused on the same 49,677 SNPs highly differentiated between the lake and stream population also underlying Fig. 6. We here performed weighted averaging of the frequency of the stream allele at each SNP between the marsh sample (M1) before the flood (top left histogram) and the lake sample closest to this marsh site (L2; bottom left). The relative weight of each sample was varied to mimic different levels of dispersal of lake fish into the marsh during the flood. We then asked what relative proportion of immigrant lake fish at the marsh site is required to yield a stream allele frequency distribution qualitatively resembling the distribution observed empirically during the flood (middle left). The evaluation of the resulting distributions (right column) was performed visually, paying particular attention to the proportion of SNPs exhibiting a stream allele frequency very near zero. This exploration revealed that the proportion of lake immigrants present at the marsh site during the flood was very high, likely around 90 - 95%.

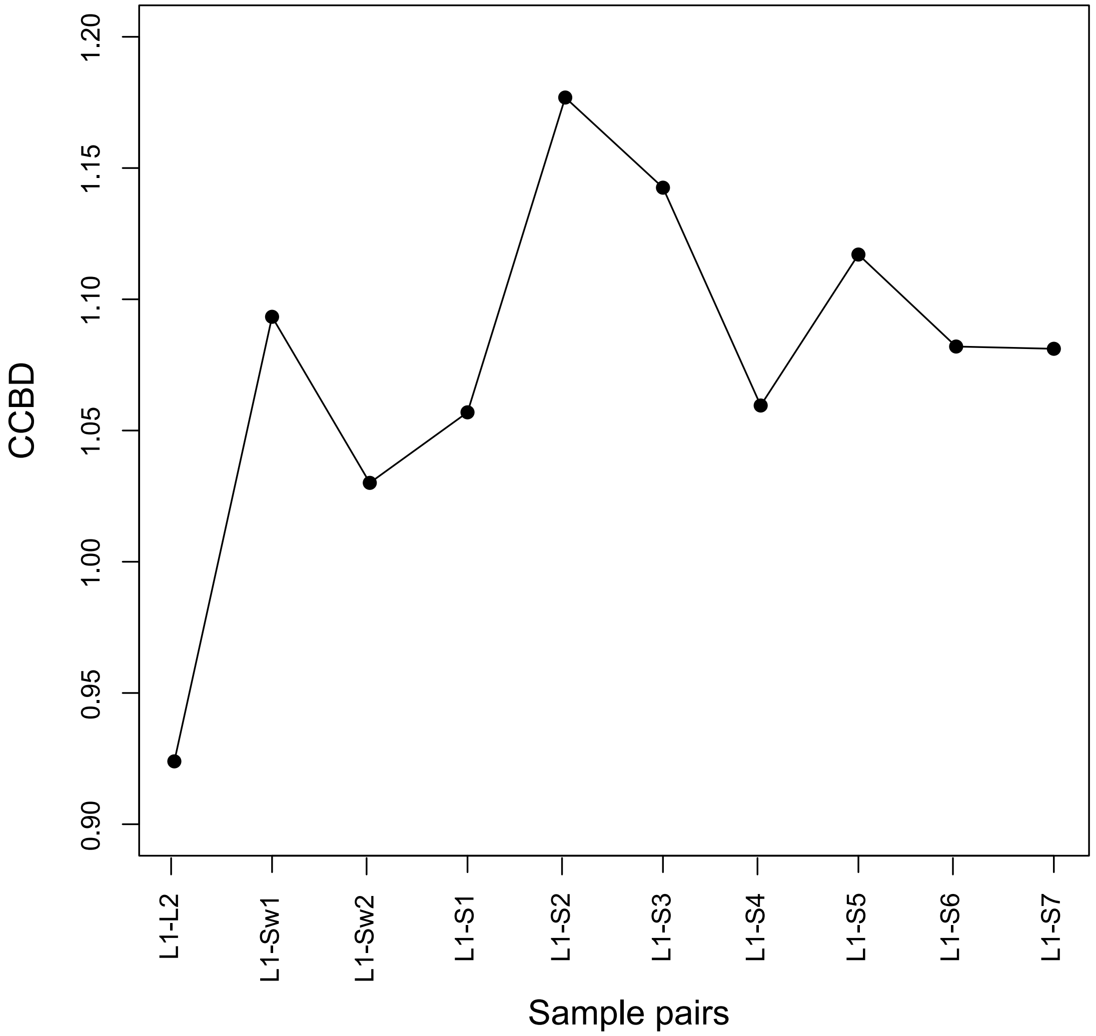

**Supplementary Figure 6**: **Alternative analysis of chromosome center-biased differentiation (CCBD).** CCBD is here calculated as described in the Methods, but the samples used for pairwise comparison were not from adjacent sites as in Fig. 2c, but involved all combinations of the samples L2-S7 with the sample L1. Consistent with Fig. 2c, the magnitude of CCBD is greatest for the L1-S2 sample pair, indicating strongest selection-gene flow antagonism in the lower reach of the inlet stream.

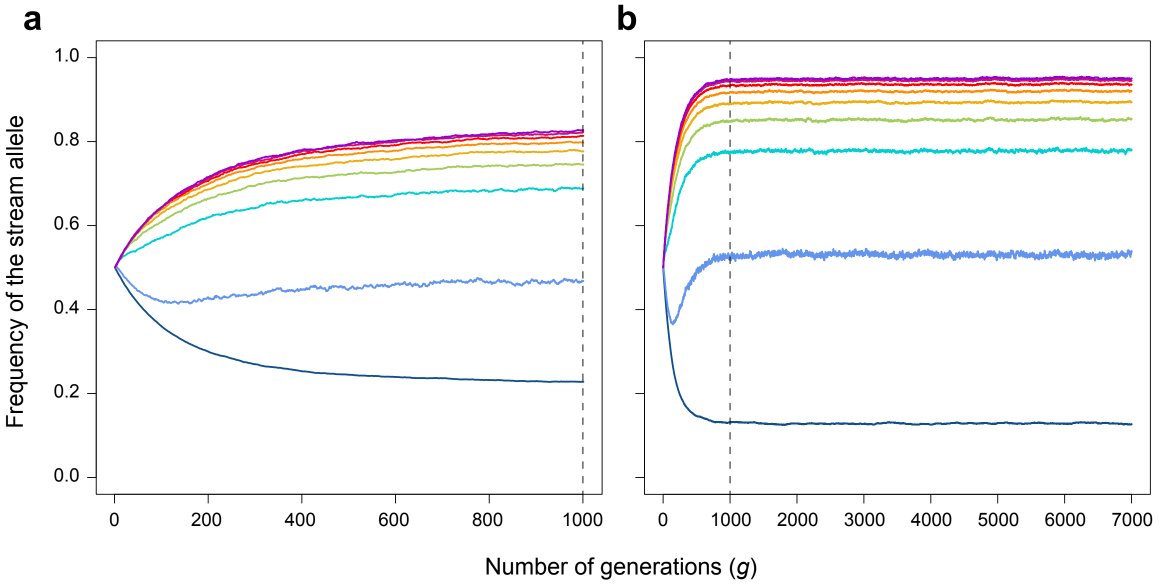

**Supplementary Figure 7**: **Determining an appropriate number of generations for the simulations.** Simulations were run with out standard stepping stone model over 1000 (a) and 7000 generations (b) for an exemplary migration rate and selection coefficient combination (*m* 0.05 and *s* 0.005). Shown is the frequency of the stream allele over time averaged over 20 simulation replications. The dotted line indicates generation 1000 in both graphs. This exploration indicates that running our simulation model over 1000 generations allows the system to approach migration-selection balance.

#### Supplementary Tables

**Supplementary Table 1**: **Characterization of the study sites in the Misty Lake watershed**. Habitat type, GPS coordinates (in decimal degrees), and the number of individuals for the genomic and morphometric analyses are given for each site. For the genomic sample sizes, the values in parentheses indicate median read depth across all genome-wide positions. The site M1 was sampled at three different time points.

| Site | Habitat | Latitude | Longitude | N genomics | N morphometrics |
|---|---|---|---|---|---|
| L1 | Lake | 50.60507824 | -127.2685989 | 62 (103) | 40 |
| L2 | Lake | 50.604347 | -127.262569 | 56 (80) | 42 |
| M1 | Marsh | 50.60516595 | -127.2579478 | 50 (133) | 36 |
| *M1 During the flood* | | | | 56 (79) | - |
| *M1 One year later* | | | | 56 (114) | - |
| M2 | Marsh | 50.605087 | -127.257812 | 56 (106) | - |
| S1 | Stream | 50.604618 | -127.257198 | 56 (74) | - |
| S2 | Stream | 50.604414 | -127.256683 | 56 (93) | - |
| S3 | Stream | 50.604375 | -127.256141 | 56 (51) | - |
| S4 | Stream | 50.603808 | -127.255397 | 40 (103) | 41 |
| S5 | Stream | 50.603056 | -127.252444 | 52 (120) | 37 |
| S6 | Stream | 50.60223555 | -127.2507798 | 56 (112) | 33 |
| S7 | Stream | 50.60060871 | -127.2476535 | 50 (107) | 41 |

**Supplementary Table 2**: **List of all genes present within a 100 kb window around each of the 34 SNPs showing complete differentiation between the most distant lake and stream sites (L1, S7).**

| SNP | Chr name | Ensembl gene ID | Gene name | Description |
|---|---|---|---|---|
| **chrI 10473231** | groupI | ENSGACG00000009957 | stx1a | syntaxin 1A (brain) [Source:ZFIN;Acc:ZDB-GENE-100802-1] |
| | groupI | ENSGACG00000009944 | SH2B2 | SH2B adaptor protein 2 [Source:HGNC Symbol;Acc:HGNC:17381] |
| | groupI | ENSGACG00000009948 | dnajc30b | DnaJ (Hsp40) homolog, subfamily C, member 30b [Source:ZFIN;Acc:ZDB-GENE-131121-532] |
| | groupI | ENSGACG00000009951 | bud23 | BUD23, rRNA methyltransferase and ribosome maturation factor [Source:ZFIN;Acc:ZDB-GENE-070410-68] |
| | groupI | ENSGACG00000009989 | caspa | caspase a [Source:ZFIN;Acc:ZDB-GENE-000616-3] |
| | groupI | ENSGACG00000009999 | rabgef1l | RAB guanine nucleotide exchange factor (GEF) 1, like [Source:ZFIN;Acc:ZDB-GENE-040426-813] |
| | groupI | ENSGACG00000010019 | NA | NA |
| | groupI | ENSGACG00000010023 | otol1b | otolin 1b [Source:ZFIN;Acc:ZDB-GENE-080416-3] |
| | groupI | ENSGACG00000010028 | nmd3 | NMD3 ribosome export adaptor [Source:ZFIN;Acc:ZDB-GENE-050320-149] |
| | groupI | ENSGACG00000010092 | NA | NULL |
| **chrI 23141976** | groupI | ENSGACG00000015056 | NA | NA |
| **chrII 13162228** | groupII | ENSGACG00000015995 | hipk3b | homeodomain interacting protein kinase 3b [Source:ZFIN;Acc:ZDB-GENE-030131-82] |
| | groupII | ENSGACG00000015997 | cstf3 | cleavage stimulation factor, 3' pre-RNA, subunit 3 [Source:ZFIN;Acc:ZDB-GENE-040426-1997] |
| | groupII | ENSGACG00000016001 | depdc7a | DEP domain containing 7, paralog a [Source:ZFIN;Acc:ZDB-GENE-060512-176] |
| | groupII | ENSGACG00000016002 | qser1 | glutamine and serine rich 1 [Source:ZFIN;Acc:ZDB-GENE-030131-2978] |
| | groupII | ENSGACG00000016006 | prrg4 | proline rich Gla (G-carboxyglutamic acid) 4 (transmembrane) [Source:ZFIN;Acc:ZDB-GENE-050417-112] |
| | groupII | ENSGACG00000016009 | eif3m | eukaryotic translation initiation factor 3, subunit M [Source:ZFIN;Acc:ZDB-GENE-040426-2643] |
| **chrIV 26355119** | groupIV | ENSGACG00000019001 | pim3 | Pim-3 proto-oncogene, serine/threonine kinase [Source:ZFIN;Acc:ZDB-GENE-050809-111] |
| | groupIV | ENSGACG00000019005 | creld2 | cysteine-rich with EGF-like domains 2 [Source:ZFIN;Acc:ZDB-GENE-040426-1626] |
| | groupIV | ENSGACG00000019007 | alg12 | ALG12, alpha-1,6-mannosyltransferase [Source:ZFIN;Acc:ZDB-GENE-041210-295] |
| | groupIV | ENSGACG00000019008 | zbed4 | zinc finger, BED-type containing 4 [Source:ZFIN;Acc:ZDB-GENE-041210-305] |
| **chrIV 26505814** | groupIV | ENSGACG00000018997 | NA | NA |
| | groupIV | ENSGACG00000018967 | NA | NA |
| | groupIV | ENSGACG00000018969 | AKR1D1 | aldo-keto reductase family 1 member D1 [Source:HGNC Symbol;Acc:HGNC:388] |
| | groupIV | ENSGACG00000018981 | kdm5a | lysine (K)-specific demethylase 5A [Source:ZFIN;Acc:ZDB-GENE-150114-1] |
| | groupIV | ENSGACG00000018983 | NA | NA |
| | groupIV | ENSGACG00000018996 | USP15 | ubiquitin specific peptidase 15 [Source:HGNC Symbol;Acc:HGNC:12613] |
| | groupIV | ENSGACG00000022283 | NA | NA |

| | | | | |
|---|---|---|---|---|
| | groupIV | ENSGACG00000018998 | IL17REL | interleukin 17 receptor E like [Source:HGNC Symbol;Acc:HGNC:33808] |
| **chrIV 27230693** | groupIV | ENSGACG00000018919 | ifrd1 | interferon-related developmental regulator 1 [Source:ZFIN;Acc:ZDB-GENE-030131-6132] |
| | groupIV | ENSGACG00000018923 | dock4b | dedicator of cytokinesis 4b [Source:ZFIN;Acc:ZDB-GENE-060130-74] |
| **chrIV 29700286** | groupIV | ENSGACG00000019720 | chchd3a | coiled-coil-helix-coiled-coil-helix domain containing 3a [Source:ZFIN;Acc:ZDB-GENE-030131-5005] |
| | groupIV | ENSGACG00000019718 | plxna4 | plexin A4 [Source:ZFIN;Acc:ZDB-GENE-030131-4663] |
| | groupIV | ENSGACG00000019721 | exoc4 | exocyst complex component 4 [Source:ZFIN;Acc:ZDB-GENE-041210-112] |
| **chrV 7120559** | groupV | ENSGACG00000003486 | mfsd13a | major facilitator superfamily domain containing 13A [Source:ZFIN;Acc:ZDB-GENE-041212-83] |
| | groupV | ENSGACG00000003470 | NA | NA |
| | groupV | ENSGACG00000003479 | CNTD1 | cyclin N-terminal domain containing 1 [Source:HGNC Symbol;Acc:HGNC:26847] |
| | groupV | ENSGACG00000003493 | sema4gb | sema domain, immunoglobulin domain (Ig), transmembrane domain (TM) and short cytoplasmic domain, (semaphorin) 4Gb [Source:ZFIN;Acc:ZDB-GENE-111117-1] |
| | groupV | ENSGACG00000003498 | mrpl43 | mitochondrial ribosomal protein L43 [Source:ZFIN;Acc:ZDB-GENE-040718-125] |
| | groupV | ENSGACG00000003504 | twnk | twinkle mtDNA helicase [Source:ZFIN;Acc:ZDB-GENE-030131-5569] |
| | groupV | ENSGACG00000003508 | lzts2b | leucine zipper, putative tumor suppressor 2b [Source:ZFIN;Acc:ZDB-GENE-030131-9587] |
| | groupV | ENSGACG00000003511 | PDZD7 | PDZ domain containing 7 [Source:HGNC Symbol;Acc:HGNC:26257] |
| | groupV | ENSGACG00000003518 | NA | NA |
| **chrVII 2342850** | groupVII | ENSGACG00000018947 | NA | NA |
| | groupVII | ENSGACG00000018955 | NA | NA |
| | groupVII | ENSGACG00000018931 | ndrg2 | NDRG family member 2 [Source:ZFIN;Acc:ZDB-GENE-041212-15] |
| | groupVII | ENSGACG00000018932 | NA | NA |
| | groupVII | ENSGACG00000018933 | si:ch211-63p21.1 | si:ch211-63p21.1 [Source:ZFIN;Acc:ZDB-GENE-141216-443] |
| | groupVII | ENSGACG00000018934 | klhl33 | kelch-like family member 33 [Source:ZFIN;Acc:ZDB-GENE-131016-4] |
| | groupVII | ENSGACG00000018940 | si:ch211-63p21.8 | si:ch211-63p21.8 [Source:ZFIN;Acc:ZDB-GENE-130530-652] |
| | groupVII | ENSGACG00000018943 | NA | NA |
| | groupVII | ENSGACG00000018948 | NA | NA |
| | groupVII | ENSGACG00000018949 | NA | NA |
| | groupVII | ENSGACG00000018954 | ltb4r | leukotriene B4 receptor [Source:ZFIN;Acc:ZDB-GENE-070705-164] |
| | groupVII | ENSGACG00000018962 | ltb4r | leukotriene B4 receptor [Source:ZFIN;Acc:ZDB-GENE-070705-164] |
| | groupVII | ENSGACG00000018963 | ltb4r2a | leukotriene B4 receptor 2a [Source:ZFIN;Acc:ZDB-GENE-140106-272] |
| | groupVII | ENSGACG00000018965 | NA | NA |
| | groupVII | ENSGACG00000018970 | NA | NA |
| **chrVIII 2559504** | groupVIII | ENSGACG00000003362 | NA | NA |
| | groupVIII | ENSGACG00000003317 | si:ch73-238c9.1 | si:ch73-238c9.1 [Source:ZFIN;Acc:ZDB-GENE-131121-81] |
| | groupVIII | ENSGACG00000003320 | reep6 | receptor accessory protein 6 [Source:ZFIN;Acc:ZDB-GENE-040912-98] |

| | | | | |
|---|---|---|---|---|
| | groupVIII | ENSGACG00000003324 | zgc:113223 | zgc:113223 [Source:ZFIN;Acc:ZDB-GENE-050320-127] |
| | groupVIII | ENSGACG00000003329 | slc1a8a | solute carrier family 1 (glutamate transporter), member 8a [Source:ZFIN;Acc:ZDB-GENE-101111-8] |
| | groupVIII | ENSGACG00000003347 | podn | podocan [Source:ZFIN;Acc:ZDB-GENE-100922-117] |
| **chrVIII 16448721** | groupVIII | ENSGACG00000011869 | mknk1 | MAP kinase interacting serine/threonine kinase 1 [Source:ZFIN;Acc:ZDB-GENE-080220-11] |
| | groupVIII | ENSGACG00000011891 | mob3c | MOB kinase activator 3C [Source:ZFIN;Acc:ZDB-GENE-040704-32] |
| | groupVIII | ENSGACG00000011897 | elovl1b | ELOVL fatty acid elongase 1b [Source:ZFIN;Acc:ZDB-GENE-040426-2755] |
| | groupVIII | ENSGACG00000011930 | NA | NA |
| | groupVIII | ENSGACG00000011937 | NA | NA |
| | groupVIII | ENSGACG00000011941 | NA | NA |
| | groupVIII | ENSGACG00000011943 | btf3l4 | basic transcription factor 3-like 4 [Source:ZFIN;Acc:ZDB-GENE-040426-1650] |
| | groupVIII | ENSGACG00000011963 | ptgfr | prostaglandin F receptor (FP) [Source:ZFIN;Acc:ZDB-GENE-120919-6] |
| | groupVIII | ENSGACG00000011969 | st6galnac5b | ST6 (alpha-N-acetyl-neuraminyl-2,3-beta-galactosyl-1,3)-N-acetylgalactosaminide alpha-2,6-sialyltransferase 5b [Source:ZFIN;Acc:ZDB-GENE-040912-25] |
| | groupVIII | ENSGACG00000011985 | slc44a5b | solute carrier family 44, member 5b [Source:ZFIN;Acc:ZDB-GENE-081105-4] |
| | groupVIII | ENSGACG00000012028 | lhx8b | LIM homeobox 8b [Source:ZFIN;Acc:ZDB-GENE-081105-153] |
| **chrX 11343215** | groupX | ENSGACG00000006652 | ppp1r9a | protein phosphatase 1, regulatory subunit 9A [Source:ZFIN;Acc:ZDB-GENE-060503-660] |
| | groupX | ENSGACG00000006481 | sp4 | sp4 transcription factor [Source:ZFIN;Acc:ZDB-GENE-031202-3] |
| | groupX | ENSGACG00000006495 | cdca7b | cell division cycle associated 7b [Source:ZFIN;Acc:ZDB-GENE-030131-607] |
| | groupX | ENSGACG00000006505 | NA | NA |
| | groupX | ENSGACG00000006508 | NA | NA |
| | groupX | ENSGACG00000006511 | col1a2 | collagen, type I, alpha 2 [Source:ZFIN;Acc:ZDB-GENE-030131-8415] |
| | groupX | ENSGACG00000006571 | casd1 | CAS1 domain containing 1 [Source:ZFIN;Acc:ZDB-GENE-060503-329] |
| | groupX | ENSGACG00000006592 | sgce | sarcoglycan, epsilon [Source:ZFIN;Acc:ZDB-GENE-030724-1] |
| **chrXIII 10658577** | groupXIII | ENSGACG00000010275 | si:dkey-40c11.2 | si:dkey-40c11.2 [Source:ZFIN;Acc:ZDB-GENE-060526-300] |
| | groupXIII | ENSGACG00000010281 | rhobtb4 | Rho related BTB domain containing 4 [Source:ZFIN;Acc:ZDB-GENE-060315-11] |
| | groupXIII | ENSGACG00000010293 | wdr54 | WD repeat domain 54 [Source:ZFIN;Acc:ZDB-GENE-040801-151] |
| | groupXIII | ENSGACG00000010296 | nol6 | nucleolar protein 6 (RNA-associated) [Source:ZFIN;Acc:ZDB-GENE-030131-6294] |
| | groupXIII | ENSGACG00000010311 | NA | NA |
| | groupXIII | ENSGACG00000010315 | aqp3a | aquaporin 3a [Source:ZFIN;Acc:ZDB-GENE-040426-2826] |
| | groupXIII | ENSGACG00000010369 | si:dkey-245n4.2 | si:dkey-245n4.2 [Source:ZFIN;Acc:ZDB-GENE-141216-258] |
| | groupXIII | ENSGACG00000010375 | NA | NA |
| **chrXVI 9022090** | groupXVI | ENSGACG00000004025 | NA | NA |
| | groupXVI | ENSGACG00000004031 | mid1 | midline 1 [Source:ZFIN;Acc:ZDB-GENE-110411-205] |
| | groupXVI | ENSGACG00000004046 | cog3 | component of oligomeric golgi complex 3 [Source:ZFIN;Acc:ZDB-GENE-050913-26] |
| | groupXVI | ENSGACG00000004048 | ednrbb | endothelin receptor type Bb [Source:ZFIN;Acc:ZDB-GENE- |

| | | | | |
|---|---|---|---|---|
| | | | | 081105-182] |
| | groupXVI | ENSGACG00000004056 | CNMD | chondromodulin [Source:HGNC Symbol;Acc:HGNC:17005] |
| | groupXVI | ENSGACG00000004057 | si:ch211-199f5.1 | protocadherin 8 [Source:HGNC Symbol;Acc:HGNC:8660] |
| **chrXVII 19022011** | groupXVII | NA | NA | NA |
| **chrXVIII 9016198** | groupXVIII | ENSGACG00000009272 | EVA1A | eva-1 homolog A, regulator of programmed cell death [Source:HGNC Symbol;Acc:HGNC:25816] |
| | groupXVIII | ENSGACG00000009276 | efhc1 | EF-hand domain (C-terminal) containing 1 [Source:ZFIN;Acc:ZDB-GENE-040426-1300] |
| | groupXVIII | ENSGACG00000009285 | tram2 | translocation associated membrane protein 2 [Source:ZFIN;Acc:ZDB-GENE-040426-2024] |
| | groupXVIII | ENSGACG00000009350 | xkr5a | XK related 5a [Source:ZFIN;Acc:ZDB-GENE-160113-27] |
| | groupXVIII | ENSGACG00000009358 | NA | NA |
| | groupXVIII | ENSGACG00000009366 | prep | prolyl endopeptidase [Source:ZFIN;Acc:ZDB-GENE-050522-14] |
| | groupXVIII | ENSGACG00000009428 | lin28b | lin-28 homolog B (C. elegans) [Source:ZFIN;Acc:ZDB-GENE-140811-1] |
| **chrXIX 8399982** | groupXIX | ENSGACG00000012555 | si:ch211-220f12.1 | FERM domain containing 4A [Source:HGNC Symbol;Acc:HGNC:25491] |
| | groupXIX | ENSGACG00000012580 | NA | NA |
| | groupXIX | ENSGACG00000012582 | impdh1a | IMP (inosine 5'-monophosphate) dehydrogenase 1a [Source:ZFIN;Acc:ZDB-GENE-040704-15] |
| | groupXIX | ENSGACG00000012608 | si:dkey-5i3.5 | si:dkey-5i3.5 [Source:ZFIN;Acc:ZDB-GENE-070424-162] |
| | groupXIX | ENSGACG00000012612 | rbm28 | RNA binding motif protein 28 [Source:ZFIN;Acc:ZDB-GENE-040426-960] |
| **chrXIX 8966355** | groupXIX | ENSGACG00000012114 | NA | NA |
| | groupXIX | ENSGACG00000012125 | psmd7 | proteasome 26S subunit, non-ATPase 7 [Source:ZFIN;Acc:ZDB-GENE-030131-5541] |
| | groupXIX | ENSGACG00000012191 | hprt1l | hypoxanthine phosphoribosyltransferase 1, like [Source:ZFIN;Acc:ZDB-GENE-040625-17] |
| | groupXIX | ENSGACG00000012213 | nudt7 | nudix (nucleoside diphosphate linked moiety X)-type motif 7 [Source:ZFIN;Acc:ZDB-GENE-131127-212] |
| | groupXIX | ENSGACG00000012216 | nfat5a | nuclear factor of activated T cells 5a [Source:ZFIN;Acc:ZDB-GENE-030131-6322] |
| | groupXIX | ENSGACG00000012221 | NA | NA |
| | groupXIX | ENSGACG00000012226 | wwox | WW domain containing oxidoreductase [Source:ZFIN;Acc:ZDB-GENE-040426-858] |
| **chrXIX 11534433** | groupXIX | ENSGACG00000010861 | si:ch211-234p18.3 | Ras protein specific guanine nucleotide releasing factor 1 [Source:HGNC Symbol;Acc:HGNC:9875] |
| | groupXIX | ENSGACG00000010863 | blm | BLM RecQ like helicase [Source:ZFIN;Acc:ZDB-GENE-070702-5] |
| | groupXIX | ENSGACG00000010873 | si:ch211-69m14.1 | NA |
| | groupXIX | ENSGACG00000010891 | malt3 | MALT paracaspase 3 [Source:ZFIN;Acc:ZDB-GENE-050419-182] |
| | groupXIX | ENSGACG00000010898 | NA | NA |
| | groupXIX | ENSGACG00000010911 | slc28a1 | solute carrier family 28 (concentrative nucleoside transporter), member 1 [Source:ZFIN;Acc:ZDB-GENE-050419-117] |
| | groupXIX | ENSGACG00000010936 | znf592 | zinc finger protein 592 [Source:ZFIN;Acc:ZDB-GENE-030131-9613] |
| **chrXIX 11670123** | groupXIX | ENSGACG00000010785 | fkbp16 | FK506 binding protein 16 [Source:ZFIN;Acc:ZDB-GENE-050208-116] |
| | groupXIX | ENSGACG00000010798 | zgc:162879 | zgc:162879 [Source:ZFIN;Acc:ZDB-GENE-070424-92] |

| | | | | |
|---|---|---|---|---|
| | groupXIX | ENSGACG00000010807 | sema4ba | sema domain, immunoglobulin domain (Ig), transmembrane domain (TM) and short cytoplasmic domain, (semaphorin) 4Ba [Source:ZFIN;Acc:ZDB-GENE-070705-31] |
| | groupXIX | ENSGACG00000022502 | NA | NA |
| | groupXIX | ENSGACG00000010823 | znf710a | zinc finger protein 710a [Source:ZFIN;Acc:ZDB-GENE-030131-5559] |
| | groupXIX | ENSGACG00000010827 | idh2 | isocitrate dehydrogenase 2 (NADP+), mitochondrial [Source:ZFIN;Acc:ZDB-GENE-031118-95] |
| **chrXIX 13388349** | groupXIX | ENSGACG00000009642 | NA | NA |
| | groupXIX | ENSGACG00000009648 | cpa4 | carboxypeptidase A4 [Source:ZFIN;Acc:ZDB-GENE-040704-61] |
| | groupXIX | ENSGACG00000009689 | smc1b | structural maintenance of chromosomes 1B [Source:ZFIN;Acc:ZDB-GENE-091217-1] |
| | groupXIX | ENSGACG00000009705 | borcs5 | BLOC-1 related complex subunit 5 [Source:ZFIN;Acc:ZDB-GENE-041010-83] |
| | groupXIX | ENSGACG00000009714 | NA | NA |
| | groupXIX | ENSGACG00000009717 | rnf141 | ring finger protein 141 [Source:ZFIN;Acc:ZDB-GENE-040625-71] |
| | groupXIX | ENSGACG00000009729 | ampd3b | adenosine monophosphate deaminase 3b [Source:ZFIN;Acc:ZDB-GENE-030131-5929] |
| | groupXIX | ENSGACG00000009748 | swap70b | switching B cell complex subunit SWAP70b [Source:ZFIN;Acc:ZDB-GENE-030131-3587] |
| **chrXIX 13544305** | groupXIX | ENSGACG00000009557 | alx1 | ALX homeobox 1 [Source:ZFIN;Acc:ZDB-GENE-050419-191] |
| | groupXIX | ENSGACG00000009562 | slc6a15 | solute carrier family 6 (neutral amino acid transporter), member 15 [Source:ZFIN;Acc:ZDB-GENE-050420-93] |
| | groupXIX | ENSGACG00000009572 | lrrk2 | leucine-rich repeat kinase 2 [Source:ZFIN;Acc:ZDB-GENE-071218-6] |
| | groupXIX | ENSGACG00000009605 | slc2a13b | solute carrier family 2 (facilitated glucose transporter), member 13b [Source:ZFIN;Acc:ZDB-GENE-090812-1] |
| | groupXIX | ENSGACG00000009605 | slc2a13b | solute carrier family 2 (facilitated glucose transporter), member 13b [Source:ZFIN;Acc:ZDB-GENE-090812-1] |
| | groupXIX | ENSGACG00000009626 | kif21a | kinesin family member 21A [Source:ZFIN;Acc:ZDB-GENE-110411-237] |
| **chrXIX 13909471** | groupXIX | ENSGACG00000009345 | si:dkey-106n21.1 | si:dkey-106n21.1 [Source:ZFIN;Acc:ZDB-GENE-131120-167] |
| | groupXIX | ENSGACG00000009373 | kitlga | kit ligand a [Source:ZFIN;Acc:ZDB-GENE-070424-1] |
| **chrXIX 14351572** | groupXIX | ENSGACG00000008891 | NA | NA |
| | groupXIX | ENSGACG00000008898 | CERK | ceramide kinase [Source:HGNC Symbol;Acc:HGNC:19256] |
| | groupXIX | ENSGACG00000008907 | NA | NA |
| | groupXIX | ENSGACG00000008914 | GRAMD4 | GRAM domain containing 4 [Source:HGNC Symbol;Acc:HGNC:29113] |
| | groupXIX | ENSGACG00000008928 | NA | NA |
| | groupXIX | ENSGACG00000008935 | drd4a | dopamine receptor D4a [Source:ZFIN;Acc:ZDB-GENE-070112-996] |
| | groupXIX | ENSGACG00000008942 | deaf1 | DEAF1 transcription factor [Source:ZFIN;Acc:ZDB-GENE-081022-163] |
| | groupXIX | ENSGACG00000008949 | BET1L | Bet1 golgi vesicular membrane trafficking protein like [Source:HGNC Symbol;Acc:HGNC:19348] |
| | groupXIX | ENSGACG00000008957 | NA | NA |
| | groupXIX | ENSGACG00000008981 | NA | NA |
| | groupXIX | ENSGACG00000008997 | NA | NA |
| | groupXIX | ENSGACG00000009014 | NA | NA |
| **chrXIX 14469026** | groupXIX | ENSGACG00000008866 | brd1b | bromodomain containing 1b [Source:ZFIN;Acc:ZDB-GENE- |

| | | | | |
|---|---|---|---|---|
| | | | | 070209-98] |
| | groupXIX | ENSGACG00000008757 | grm8b | glutamate receptor, metabotropic 8b [Source:ZFIN;Acc:ZDB-GENE-110421-3] |
| | groupXIX | ENSGACG00000008779 | NA | NA |
| | groupXIX | ENSGACG00000008783 | NA | NA |
| | groupXIX | ENSGACG00000008835 | si:ch211-253p14.2 | si:ch211-253p14.2 [Source:ZFIN;Acc:ZDB-GENE-131127-524] |
| | groupXIX | ENSGACG00000008837 | NA | NA |
| | groupXIX | ENSGACG00000008843 | ftsj1 | FtsJ RNA methyltransferase homolog 1 [Source:ZFIN;Acc:ZDB-GENE-041114-83] |
| | groupXIX | ENSGACG00000008869 | NA | NA |
| | groupXIX | ENSGACG00000008872 | NA | NA |
| | groupXIX | ENSGACG00000008877 | NA | NA |
| | groupXIX | ENSGACG00000008880 | TAFA5 | TAFA chemokine like family member 5 [Source:HGNC Symbol;Acc:HGNC:21592] |
| **chrXIX 14814192** | groupXIX | ENSGACG00000008655 | CADPS2 | calcium dependent secretion activator 2 [Source:HGNC Symbol;Acc:HGNC:16018] |
| | groupXIX | ENSGACG00000008606 | NA | NA |
| | groupXIX | ENSGACG00000008612 | NA | NA |
| | groupXIX | ENSGACG00000008617 | NA | NA |
| | groupXIX | ENSGACG00000008638 | aass | aminoadipate-semialdehyde synthase [Source:ZFIN;Acc:ZDB-GENE-061220-8] |
| | groupXIX | ENSGACG00000008652 | fezf1 | FEZ family zinc finger 1 [Source:ZFIN;Acc:ZDB-GENE-060929-970] |
| **chrXIX 14917217** | groupXIX | ENSGACG00000008512 | NA | NA |
| | groupXIX | ENSGACG00000008520 | glg1a | golgi glycoprotein 1a [Source:ZFIN;Acc:ZDB-GENE-030131-6448] |
| | groupXIX | ENSGACG00000008531 | rfwd3 | ring finger and WD repeat domain 3 [Source:ZFIN;Acc:ZDB-GENE-120529-1] |
| | groupXIX | ENSGACG00000008536 | MLKL | mixed lineage kinase domain like pseudokinase [Source:HGNC Symbol;Acc:HGNC:26617] |
| | groupXIX | ENSGACG00000008543 | NA | NA |
| | groupXIX | ENSGACG00000008550 | NA | NA |
| | groupXIX | ENSGACG00000008572 | NA | NA |
| | groupXIX | ENSGACG00000008577 | NA | NA |
| | groupXIX | ENSGACG00000008587 | NA | NA |
| | groupXIX | ENSGACG00000008589 | FAAP24 | FA core complex associated protein 24 [Source:HGNC Symbol;Acc:HGNC:28467] |
| | groupXIX | ENSGACG00000008594 | NA | NA |
| | groupXIX | ENSGACG00000008599 | faap24 | FA core complex associated protein 24 [Source:ZFIN;Acc:ZDB-GENE-070410-53] |
| | groupXIX | ENSGACG00000008606 | NA | NA |
| **chrXIX 15057480** | groupXIX | ENSGACG00000008453 | cacna2d4b | calcium channel, voltage-dependent, alpha 2/delta subunit 4b [Source:ZFIN;Acc:ZDB-GENE-100422-18] |
| | groupXIX | ENSGACG00000008376 | lsp1 | lymphocyte specific protein 1 [Source:ZFIN;Acc:ZDB-GENE-131127-171] |
| | groupXIX | ENSGACG00000008376 | lsp1 | lymphocyte specific protein 1 [Source:ZFIN;Acc:ZDB-GENE-131127-171] |
| | groupXIX | ENSGACG00000008384 | TNNT3 | troponin T3, fast skeletal type [Source:HGNC Symbol;Acc:HGNC:11950] |
| | groupXIX | ENSGACG00000008444 | myod1 | myogenic differentiation 1 [Source:ZFIN;Acc:ZDB-GENE-980526-561] |

| | | | | |
|---|---|---|---|---|
| | groupXIX | ENSGACG00000008450 | NA | NA |
| | groupXIX | ENSGACG00000008453 | cacna2d4b | calcium channel, voltage-dependent, alpha 2/delta subunit 4b [Source:ZFIN;Acc:ZDB-GENE-100422-18] |
| **chrXIX 15702851** | groupXIX | ENSGACG00000007601 | PTPRB | protein tyrosine phosphatase receptor type B [Source:HGNC Symbol;Acc:HGNC:9665] |
| | groupXIX | ENSGACG00000007499 | NA | NA |
| | groupXIX | ENSGACG00000007525 | lamb1a | laminin, beta 1a [Source:ZFIN;Acc:ZDB-GENE-021226-1] |
| | groupXIX | ENSGACG00000007545 | NA | NA |
| | groupXIX | ENSGACG00000007560 | nup160 | nucleoporin 160 [Source:ZFIN;Acc:ZDB-GENE-040426-1603] |
| | groupXIX | ENSGACG00000007588 | frs2b | fibroblast growth factor receptor substrate 2b [Source:ZFIN;Acc:ZDB-GENE-040718-406] |
| **chrXX 3378337** | groupXX | ENSGACG00000013330 | tspan13a | tetraspanin 13a [Source:ZFIN;Acc:ZDB-GENE-040718-19] |
| | groupXX | ENSGACG00000013308 | pear1 | platelet endothelial aggregation receptor 1 [Source:ZFIN;Acc:ZDB-GENE-091230-1] |
| | groupXX | ENSGACG00000013318 | gpatch4 | G patch domain containing 4 [Source:ZFIN;Acc:ZDB-GENE-041008-158] |
| | groupXX | ENSGACG00000013326 | NA | NA |
| | groupXX | ENSGACG00000013327 | NA | NA |
| | groupXX | ENSGACG00000013337 | bag6 | BCL2 associated athanogene 6 [Source:ZFIN;Acc:ZDB-GENE-010501-5] |
| | groupXX | ENSGACG00000013362 | NA | NA |
| | groupXX | ENSGACG00000013372 | tnfb | tumor necrosis factor b (TNF superfamily, member 2) [Source:ZFIN;Acc:ZDB-GENE-050601-2] |
| | groupXX | ENSGACG00000013381 | GABBR1 | gamma-aminobutyric acid type B receptor subunit 1 [Source:HGNC Symbol;Acc:HGNC:4070] |
| **chrXXI 9639790** | groupXXI | ENSGACG00000002461 | CNTNAP2 | contactin associated protein like 2 [Source:HGNC Symbol;Acc:HGNC:13830] |
| | groupXXI | ENSGACG00000002473 | NA | NA |
| **chrXXI 10675081** | groupXXI | ENSGACG00000002744 | jph1a | junctophilin 1a [Source:ZFIN;Acc:ZDB-GENE-040724-233] |
| | groupXXI | ENSGACG00000002714 | crispld1a | cysteine-rich secretory protein LCCL domain containing 1a [Source:ZFIN;Acc:ZDB-GENE-090612-1] |
| | groupXXI | ENSGACG00000002723 | pi15a | peptidase inhibitor 15a [Source:ZFIN;Acc:ZDB-GENE-040724-135] |
| | groupXXI | ENSGACG00000002727 | gdap1 | ganglioside induced differentiation associated protein 1 [Source:ZFIN;Acc:ZDB-GENE-050522-424] |
| | groupXXI | ENSGACG00000002748 | eloca | elongin C paralog a [Source:ZFIN;Acc:ZDB-GENE-040912-120] |
| | groupXXI | ENSGACG00000002752 | flj11011l | hypothetical protein FLJ11011-like (H. sapiens) [Source:ZFIN;Acc:ZDB-GENE-050113-1] |
| | groupXXI | ENSGACG00000002757 | stau2 | staufen double-stranded RNA binding protein 2 [Source:ZFIN;Acc:ZDB-GENE-040426-687] |
| **chrXXI 12602955** | groupXXI | ENSGACG00000003687 | si:ch73-206p6.1 | si:ch73-206p6.1 [Source:ZFIN;Acc:ZDB-GENE-030131-5048] |
| | groupXXI | ENSGACG00000003693 | plod2 | procollagen-lysine, 2-oxoglutarate 5-dioxygenase 2 [Source:ZFIN;Acc:ZDB-GENE-070326-1] |
| **chrXXI 12862781** | groupXXI | ENSGACG00000003756 | pcolce2b | procollagen C-endopeptidase enhancer 2b [Source:ZFIN;Acc:ZDB-GENE-040426-1177] |
| | groupXXI | ENSGACG00000003763 | u2surp | U2 snRNP-associated SURP domain containing [Source:ZFIN;Acc:ZDB-GENE-070912-400] |
| | groupXXI | ENSGACG00000003774 | acad11 | acyl-CoA dehydrogenase family, member 11 [Source:ZFIN;Acc:ZDB-GENE-040426-814] |
| | groupXXI | ENSGACG00000003814 | ackr4b | atypical chemokine receptor 4b [Source:ZFIN;Acc:ZDB- |

| | | | | |
|---|---|---|---|---|
| | | | | GENE-051107-10] |
| | groupXXI | ENSGACG00000003816 | uba5 | ubiquitin-like modifier activating enzyme 5 [Source:ZFIN;Acc:ZDB-GENE-031112-2] |
| | groupXXI | ENSGACG00000003823 | nphp3 | nephronophthisis 3 [Source:ZFIN;Acc:ZDB-GENE-091204-117] |
| | groupXXI | ENSGACG00000003837 | si:dkey-96n2.3 | si:dkey-96n2.3 [Source:ZFIN;Acc:ZDB-GENE-091204-118] |
| | groupXXI | ENSGACG00000003838 | si:ch211-285f17.1 | si:ch211-285f17.1 [Source:ZFIN;Acc:ZDB-GENE-000607-77] |
